# The Gibbs free energy landscape based on liver metabolome revealed thermodynamic robustness against fasting and obesity

**DOI:** 10.64898/2025.12.04.692290

**Authors:** Takumi Abekawa, Satoshi Ohno, Akiyoshi Hirayama, Tomoyoshi Soga, Shinya Kuroda

## Abstract

Mammalian liver metabolism undergoes a substantial shift during fasting. Thermodynamic principles impose fundamental constraints on metabolism, and the Gibbs free energy change of reaction (Δ_*r*_*G*′) indicates the reaction’s direction and distance from equilibrium. However, Δ_*r*_*G*′ landscapes in intact mammalian organs remain largely uncharacterized. Here, we mapped Δ_*r*_*G*′ profile of glucose metabolism in mouse liver during fasting, using experimentally measured absolute metabolite concentrations and a newly developed computational method, GLEAM. We found that despite large metabolite fluctuations during fasting, the corresponding Δ_*r*_*G*′s remained robust, even for reactions reversing the direction between glycolysis and gluconeogenesis. A remarkable thermodynamic robustness is found in maintaining potential candidates of rate-limiting steps during fasting. This robustness is achieved by expending substantial costs for enzyme expressions, contributing to efficient switching from glycolysis to gluconeogenesis. Furthermore, obese mouse liver also showed thermodynamic robustness despite obesity-induced metabolic disruption. Our framework provided a novel thermodynamic perspective on intact organ metabolism, demonstrating that the liver robustly maintains thermodynamic characteristics favorable for metabolic control by buffering metabolite concentration differences.

## INTRODUCTION

Blood glucose homeostasis is crucial in mammals for providing stable energy sources to organs and maintaining health. During fasting, the liver plays a central role in maintaining blood glucose homeostasis by shifting its metabolism from glycolysis to gluconeogenesis ^1–5^. This metabolic shift is a highly regulated process that involves the directional reversal of metabolic reactions and fluctuations in the concentrations of numerous metabolites ^6–8^. Obesity induces metabolic dysregulation in the liver ^7–12^, leading to impaired glucose homeostasis ^13–17^ and the development of further diseases, including type 2 diabetes ^9,18–20^.

Metabolic processes are governed by the laws of thermodynamics. According to the second law of thermodynamics, in the cellular environment, metabolic reactions proceed in the direction where the Gibbs free energy change of reaction (Δ_*r*_*G*′) is negative, and Δ_*r*_*G*′ is zero when the reaction reaches equilibrium. Therefore, Δ_*r*_*G*′ indicates the reaction distance from equilibrium. Δ_*r*_*G*′ can be calculated as the difference between the Gibbs free energy of formation (Δ_*f*_*G*′) of the product and substrate metabolites:

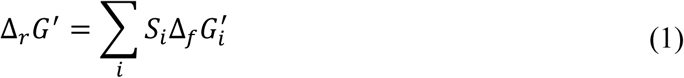

where *S*_*i*_ is the stoichiometric coefficient of metabolite *i* in the reaction. Δ_*f*_*G*′ depends on the metabolite activity, practically substituted by the concentration:

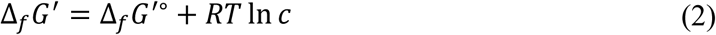

where Δ_*f*_*G*′∘ is the standard Gibbs free energy of formation representing Δ_*f*_*G*′ at 1 M of metabolite concentration, *R* is the gas constant, *T* is temperature, and *c* is metabolite concentration. The prime placed in Δ_*f*_*G*′∘, Δ_*f*_*G*′, and Δ_*r*_*G*′ denotes that those values incorporate the effect of a particular pH, pMg, and ionic strength on metabolite activity. Although calculation of Δ_*r*_*G*′requires absolute metabolite concentrations, mass spectrometry-based metabolomics analyses often suffer from ion suppression, a phenomenon where co-eluting compounds compromise quantitative accuracy by reducing analyte ionization efficiency ^21^. Nevertheless, previous studies have successfully achieved absolute quantification of intracellular metabolites, primarily by employing isotope-labeled internal standards ^22,23^. These techniques have enabled the investigation of absolute metabolite concentrations and subsequent thermodynamic analyses, such as Δ_*r*_*G*′ landscapes, in microorganisms and cultured cells. ^24–27^. Despite these advancements, thermodynamic profiles have rarely been investigated at *in vivo* mammalian organ levels. Furthermore, all prior thermodynamic studies have focused on glycolysis-dominant conditions where glucose is the primary nutrient, leaving a large gap in our understanding of metabolism during fasting, a condition dominated by gluconeogenesis.

Mathematical metabolic modeling is an essential framework for a quantitative system-level understanding of metabolism. The thermodynamic principle regarding reaction direction has been widely applied as a constraint to increase the reliability of metabolic models ^28–35^. Furthermore, the significance of Δ_*r*_*G*′ extends beyond physical aspects such as distance from equilibrium and reaction directionality to several physiological meanings, including pathway rate-limiting steps quantified by the flux control coefficient (FCC) ^36–38^, and enzyme demands for carrying metabolic flux ^39–42^. However, these thermodynamic frameworks have hardly been applied to experimental data, especially those from multicellular organisms.

In this study, we explored Δ_*r*_*G*′ landscape in the liver metabolism of mice at different fasting levels using time-series absolute quantitative metabolomic data measured by CE-MS, which can reduce the effect of ion suppression ^43,44^, without using isotopically labeled standards. Our results are the first *in vivo* identification of large-scale Δ_*r*_*G*′ landscapes in mammalian organisms, and also the first elucidation of Δ_*r*_*G*′ behavior during fasting in any cell types. Since metabolomic data contains uncertainties, we developed GLEAM (Gibbs free energy of reaction Landscape Estimation from metabolome Assisted by variance-covariance Matrix) to estimate Δ_*r*_*G*′ with thermodynamically consistent metabolite concentrations from experimental metabolomic data. The estimated Δ_*r*_*G*′ showed that the global Δ_*r*_*G*′landscape in mouse liver is robust during fasting, despite large fluctuations in metabolite concentrations. We further integrated the Δ_*r*_*G*′ and enzyme kinetics to investigate the physiological significance of the Δ_*r*_*G*′ landscape in terms of rate-limiting steps and efficiency of metabolic switching. In addition to the thermodynamic robustness of mouse liver against fasting, we also demonstrated the robustness against obesity using leptin-deficient obese (*ob*/*ob*) mice ^45,46^. Thus, our study provides novel insights into glucose metabolism in *in vivo* mammalian liver from a thermodynamic standpoint. These findings extend our understanding of mammalian metabolisms beyond individual metabolite concentrations.

## RESULTS

### Δ_*r*_*G*′ landscape is robust against fasting despite metabolite concentration fluctuations

In this study, we developed GLEAM to estimate Δ_*r*_*G*′ from time-series metabolome and Δ_*f*_*G*′∘. One of the challenges in estimating Δ_*r*_*G*′ in an entire reaction pathway is inherent uncertainties and missing values in metabolome and Δ_*f*_*G*′∘ data. To address this issue, GLEAM utilizes thermodynamic constraints and variances and covariances of metabolome and Δ_*f*_*G*′∘ data, and estimates a complete set of thermodynamically consistent metabolite concentrations and Δ_*f*_*G*′∘, and subsequent Δ_*r*_*G*′ (see Methods). Briefly, GLEAM fits metabolite concentrations and Δ_*f*_*G*′∘ to data from the experiment or database within a range that every reaction holds Δ_*r*_*G*′ < 0 by the weighted least mean squares method. The weights of the residuals are determined by variances and covariances of metabolite concentrations and Δ_*f*_*G*′∘ data.

By applying GLEAM to time-series metabolomic data from *ad libitum* feeding to fasting in mice, and Δ_*f*_*G*′∘ data from eQuilibrator (Table S1), we estimated the Δ_*r*_*G*′ landscape (Fig. 1A, B, Table S2) in glycolysis, gluconeogenesis, and TCA cycle (Table S3) in the liver. At *ad libitum* feeding (0-hour fasting), when glycolysis is dominant, most of the irreversible reactions exhibited Δ_*r*_*G*′ < −10 kJ/mol, which were defined as far-from-equilibrium reactions, such as Δ_*r*_*G*′ = −21.56 kJ/mol in PFK and Δ_*r*_*G*′ = −34.34 kJ/mol in PDH. Among the 11 irreversible reactions, only three reactions (ACO:IDH, PC, and PCK) exhibited Δ_*r*_*G*′ > −10 kJ/mol, and PC and PCK were exceptionally close to equilibrium, with Δ_*r*_*G*′ = −3.48 kJ/mol and −3.70 kJ/mol, respectively. In contrast, most of the reversible reactions exhibited Δ_*r*_*G*′ > −5 kJ/mol, which were defined as near-equilibrium reactions, at *ad libitum* feeding, such as Δ_*r*_*G*′ = −0.24 kJ/mol in GPI and Δ_*r*_*G*′ = −0.03 kJ/mol in SDH. Four reversible reactions (ALDO, LSC, MDH, and ME) were exceptions that exhibited Δ_*r*_*G*′ < −5 kJ/mol. Among those reactions, MDH was largely away from equilibrium in the opposite direction of the oxidative TCA cycle, with Δ_*r*_*G*′ = −18.13 kJ/mol.

**Fig. 1:**
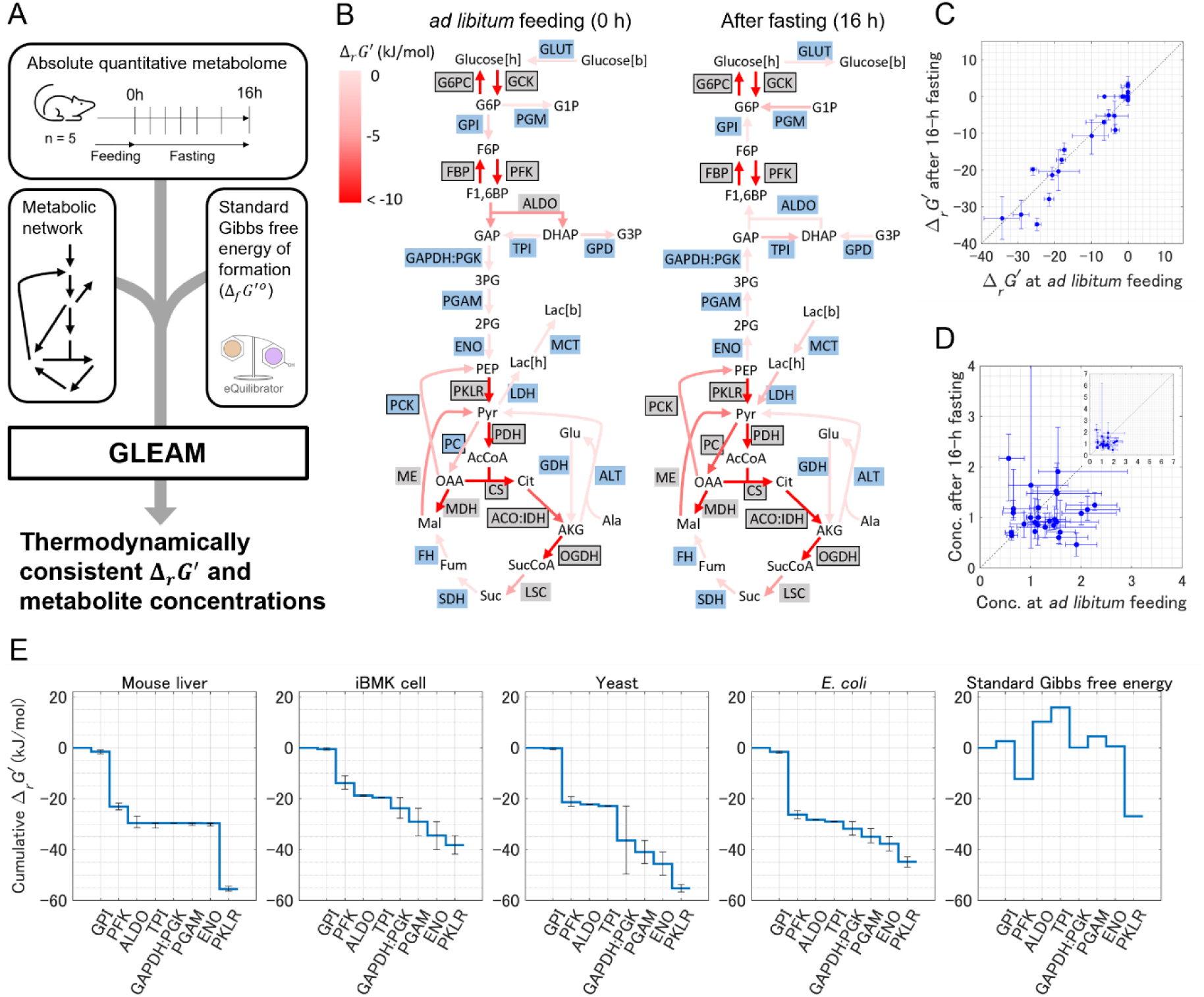
Δ_*r*_*G*′ landscape is robust against fasting despite metabolite concentration fluctuations. (**A**) Overview of the application of GLEAM to estimate Δ_*r*_*G*′ landscape in the mouse liver. Metabolomic data were acquired from the liver, skeletal muscle, and plasma of WT and *ob*/*ob* mice during fasting (Table S1) ^8,66^. *n* = 5 biological replicates were used per group. The estimated metabolite concentrations, Δ_*f*_*G*′∘, and Δ_*r*_*G*′ values by GLEAM are shown in Table S2. (**B**) Δ_*r*_*G*′ landscape of glucose metabolism in mouse liver at *ad libitum* feeding and after 16-hour fasting. Δ_*r*_*G*′ is depicted by arrow colors. Shaded texts represent reaction names; boxed reactions are irreversible, while unboxed reactions are reversible. Blue shadings indicate near-equilibrium reactions with Δ_*r*_*G*′ > −5 kJ/mol, and gray shadings indicate the other reactions with Δ_*r*_*G*′ ≤ −5 kJ/mol. Unshaded texts represent metabolites. [h], [m], and [b] indicate hepatic, myocytic, and blood metabolites, respectively. Abbreviations of reactions and metabolites are defined in the glossary and Table S3. (**C**) Comparison of Δ_*r*_*G*′ in WT mouse liver at *ad libitum* feeding and after 16-hour fasting. The forward direction (Δ_*r*_*G*′ < 0) is defined as the reaction direction at *ad libitum* feeding. Error bars show 95% confidence intervals. (**D**) Comparison of metabolite concentrations in WT mouse liver at *ad libitum* feeding and after 16-hour fasting. Plotted metabolite concentrations represent those normalized by the mean value at all seven time points and two genotypes (14 conditions in total) to eliminate scale differences in concentrations among metabolites. Error bars show 95% confidence intervals. (**E**) Comparison of Δ_*r*_*G*′ in glycolysis across cell types and organisms. The data in iBMK cell, yeast, and *E. coli* were measured using LC-MS and isotope tracers ^26^. Standard Gibbs free energy was calculated by treating all metabolite concentrations as 1 M. The vertical axis represents cumulative Δ_*r*_*G*′ from GPI. Error bars show 95% confidence intervals.

The fasting-induced metabolic shift in the liver from glycolysis to gluconeogenesis may lead to changes in thermodynamic characteristics of metabolic reactions, including reversal of the sign of Δ_*r*_*G*′ of some reactions *i.e.,* reversal of their direction, by fluctuations in several metabolite concentrations. After 16-hour fasting when gluconeogenesis is dominant, Δ_*r*_*G*′values remained similar in the liver (Fig. 1C), with variation in 24/29 reactions being less than 5 kJ/mol. As a result of this thermodynamic robustness, all irreversible reactions exhibited Δ_*r*_*G*′ < −10 kJ/mol after 16-hour fasting, indicating that the overall trend of irreversible reactions being far-from-equilibrium reactions was maintained across *ad libitum* feeding and after fasting. Although PC and PCK had been near-equilibrium at *ad libitum* feeding, these reactions moved away from equilibrium to exhibit Δ_*r*_*G*′ = −9.07 kJ/mol and −5.25 kJ/mol after fasting, respectively. As for reversible reactions, despite reversal of some directions to switch the liver’s metabolism from glycolysis to gluconeogenesis, all the reversible reactions with Δ_*r*_*G*′ > −5 kJ/mol at *ad libitum* feeding remained Δ_*r*_*G*′ > −5 kJ/mol, indicating that the overall trend of reversible reactions being near-equilibrium reactions was maintained across *ad libitum* feeding and after fasting. In contrast to Δ_*r*_*G*′, individual metabolite concentrations widely fluctuated during 16-hour fasting (Fig. 1D), with log2 fold change exceeding 0.5 for over 50% of metabolites. This result suggests that the thermodynamic architecture in glucose metabolism was robust against fasting while metabolite concentrations were reorganized. Note that the robustness of Δ_*r*_*G*′ and metabolite concentrations cannot be directly compared by correlation coefficients for Fig. 1C and D because Δ_*r*_*G*′ values were not normalized while metabolite concentrations were normalized, leading to much higher correlation coefficient for Δ_*r*_*G*′. The reason why Δ_*r*_*G*′values were not normalized in Fig. 1C is that Δ_*r*_*G*′ does not represent abundance and thus should be evaluated their robustness not by fold changes but by difference values.

To examine whether the thermodynamic characteristics of metabolic reactions observed in mouse liver are conserved across species, we compared the estimated Δ_*r*_*G*′ in glycolytic reactions in mouse liver with those in other biological systems. The previous study ^26^ determined Δ_*r*_*G*′ in glycolysis of immortalized baby mouse kidney epithelial (iBMK) cells, yeast, and *Escherichia coli*, reporting that in all these cell types, similar to mouse liver, reversible reactions were closer to equilibrium compared to the irreversible reaction, PFK (Fig. 1E). This result suggests that the thermodynamic characteristics in glycolysis are widely conserved across species. An exception is PKLR, whose Δ_*r*_*G*′ in mouse liver was more than 20 kJ/mol smaller than those in the other cell types. Among the metabolite concentrations (or ratios) involved in PKLR, pyruvate concentration and ATP/ADP were smaller in mouse liver than in mammalian iBMK cells, yeast, and *E. coli* (Table S4). In particular, pyruvate concentrations were two orders of magnitude smaller in mouse liver (3.77 × 10^-2^ mM) compared to the other three cell types (5.88 mM in iBMK cells, 9.40 mM in yeast, and 3.66 mM in *E. coli*).

### Δ_*r*_*G*′ maintenance during fasting is associated with metabolite concentration fluctuations in different ways among pathways

To characterize the metabolic reactions exhibiting large thermodynamic changes accompanying the fasting-induced metabolic shift, we linked the attributes of reactions including reversibility and pathway to Δ_*r*_*G*′_16h_− Δ_*r*_*G*′_0h_, defined as Δ_*r*_*G*′ variations during fasting (Fig. 2A). Among the seven irreversible reactions of glycolysis and gluconeogenesis, four reactions (GCK, PFK, PKLR, and PC) exhibited an absolute Δ_*r*_*G*′ variation during fasting greater than 5 kJ/mol. In contrast, only one reversible reaction (ALDO) out of 11 showed an absolute Δ_*r*_*G*′ variation during fasting exceeding 5 kJ/mol, despite the reversal of reversible reaction directions between glycolysis and gluconeogenesis under these conditions. This result suggests that reversible reactions maintained Δ_*r*_*G*′ more strictly than irreversible reactions even though the sign of Δ_*r*_*G*′ is reversed. In the TCA cycle, the absolute Δ_*r*_*G*′ variations during fasting were less than 3 kJ/mol for all reactions, indicating that the TCA cycle is thermodynamically stable in response to fasting.

**Fig. 2:**
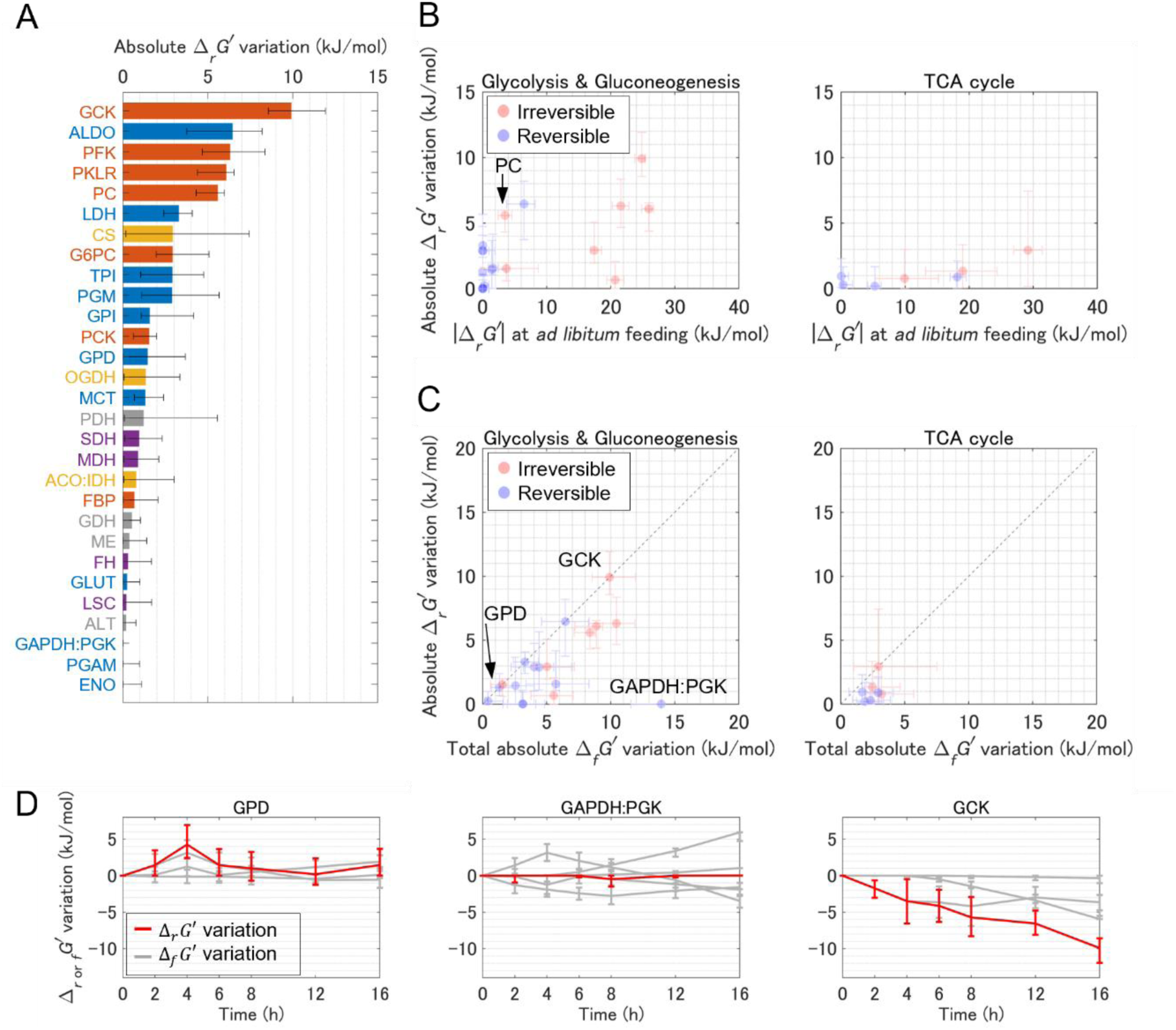
Δ_*r*_*G*′ maintenance during fasting is associated with metabolite concentration fluctuations in different ways among pathways. (**A**) Absolute Δ_*r*_*G*′ variation = |Δ_*r*_*G*′_16h_− Δ_*r*_*G*′_0h_|. Bars depict irreversible reactions in glycolysis and gluconeogenesis (orange), reversible reactions in glycolysis and gluconeogenesis (blue), irreversible reactions in the TCA cycle (yellow), reversible reactions in the TCA cycle (purple), and the other reactions in the metabolic network (grey), respectively. Error bars show 95% confidence intervals. (**B**) Relationship between absolute Δ_*r*_*G*′ variation and reaction distance from equilibrium represented by absolute Δ_*r*_*G*′ at *ad libitum* feeding. Error bars show 95% confidence intervals. The reaction PC is highlighted and discussed in the Results section. (**C**) Comparison of absolute Δ_*r*_*G*′ variation of reactions and total absolute Δ_*f*_*G*′ variation of metabolites. Absolute Δ_*f*_*G*′ variation is dependent on concentrations of metabolites involved in the reaction, and defined by ∑|*S*(Δ_*f*_*G*′ − Δ_*f*_*G*′_0*h*_)|. When Δ_*f*_*G*′ variation = *S*(Δ_*f*_*G*′_16*h*_− Δ_*f*_*G*′_0*h*_) of the metabolites involved in the reaction are cancelled out, absolute Δ_*r*_*G*′ variation is smaller than total absolute Δ_*f*_*G*′variation, resulting in the reaction being plotted in the lower right. Plotted reactions for glycolysis and gluconeogenesis, and the TCA cycles are the same as Fig. 2B. Error bars show 95% confidence intervals. The reactions GPD, GAPDH:PGK and GCK are highlighted and discussed in the Results section. (**D**) Time courses of Δ_*r*_*G*′ variation and Δ_*f*_*G*′ variation for representative reactions: GPD (small Δ_*f*_*G*′ variations), GAPDH:PGK (large Δ_*f*_*G*′variations which are canceled out), and GCK (large Δ_*f*_*G*′ variations which are not canceled out). Red and grey lines depict Δ_*r*_*G*′ variation and Δ_*f*_*G*′ variation, respectively. Error bars show 95% confidence intervals.

As Δ_*r*_*G*′ variations during fasting differed between irreversible and reversible reactions in glycolysis and gluconeogenesis, we further examined the relationship between the distance from equilibrium (|Δ_*r*_*G*′|) at *ad libitum* feeding and the absolute Δ_*r*_*G*′ variation (Fig. 2B). In glycolysis and gluconeogenesis, the absolute Δ_*r*_*G*′ variations during fasting were less than 3.5 kJ/mol for near-equilibrium reactions with |Δ_*r*_*G*′| < 5 kJ/mol at *ad libitum* feeding except for PC. PC moved away from equilibrium only when gluconeogenesis is dominant in response to fasting. In contrast, the absolute Δ_*r*_*G*′ variations of the reactions with |Δ_*r*_*G*′| ≥ 5 kJ/mol at *ad libitum* feeding ranged between 0 and 10 kJ/mol, and were significantly larger than those of the near-equilibrium reactions (p value = 0.037 < 0.05 calculated by two-tailed Welch’s *t*-test, n = 12 for near-equilibrium reactions with |Δ_*r*_*G*′| < 5 and n = 6 for the other reactions with |Δ_*r*_*G*′ | ≥ 5 kJ/mol). This result suggests that Δ_*r*_*G*′ of near-equilibrium reactions in glycolysis and gluconeogenesis tended to be strictly maintained, while this tendency was lost among other reactions. In the TCA cycle, Δ_*r*_*G*′ was strictly maintained regardless of the distance from equilibrium. The statistical test was not performed for the TCA cycle due to a lack of near-equilibrium reactions.

To investigate the reason maintaining Δ_*r*_*G*′ in near-equilibrium reactions in glycolysis and gluconeogenesis, as well as in the TCA cycle reactions, we compared the absolute Δ_*r*_*G*′variation with the total absolute Δ_*f*_*G*′ variation of metabolites involved in the reaction (Fig. 2C). The total absolute Δ_*f*_*G*′ variation of metabolites is defined as ∑|*S*(Δ_*f*_*G*′_16h_ − Δ_*f*_*G*′_0h_)|, where *S* is the stoichiometric coefficient of a metabolite involved in the reaction. The Δ_*r*_*G*′variation is associated in various ways with fluctuations in the Δ_*f*_*G*′ of substrates and products, which depend on the metabolite concentrations. For example, when substrate and product concentrations remain relatively constant and changes in their Δ_*f*_*G*′ remain small, the Δ_*r*_*G*′ variation is small (*e.g.*, GPD in Fig. 2D). Even when substrate and product concentrations largely fluctuate and changes in their Δ_*f*_*G*′are large, the Δ_*r*_*G*′ variation can remain small when the changes in Δ_*f*_*G*′ of substrate and products are cancelled out. This can happen, for instance, when both the substrate and product Δ_*f*_*G*′ values increase simultaneously (*e.g.*, GAPDH:PGK in Fig. 2D). In contrast, when the changes in Δ_*f*_*G*′ of substrate and products are not cancelled out, the Δ_*r*_*G*′ variation becomes large (*e.g.*, GCK in Fig. 2D). In glycolysis and gluconeogenesis, reactions with small absolute Δ_*r*_*G*′ variation due to small fluctuation in metabolite concentrations, such as GPD, and due to the cancellation of Δ_*f*_*G*′ variations such as GAPDH:PGK, were both observed (Fig. 2C). The existence of the latter type of reaction suggests that while metabolite concentrations fluctuate in response to fasting in glycolysis and gluconeogenesis, the fluctuations in metabolites involved in near-equilibrium reactions were coordinated, thereby maintaining Δ_*r*_*G*′ of these reactions. In the TCA cycle, the reason for maintaining the absolute Δ_*r*_*G*′ variations were small fluctuations in metabolite concentrations for all the reactions.

### Thermodynamic Rate-limiting Candidates (TRCs) were maintained during fasting

One of the physiological consequences of reaction distances from equilibrium is the control of steady-state pathway fluxes. The extent to which perturbation of each reaction can control pathway steady-state flux is quantified by the flux control coefficient (FCC) in metabolic control analysis (MCA) ^36,47^. In this study, we analytically calculated FCCs under both arbitrary Δ_*r*_*G*′ landscapes and those observed in mouse liver within glycolysis (Fig. 3A), gluconeogenesis (Fig. 3B), and the TCA cycle (Fig. 3C) from an enzymatic rate law using the MCA’s theorems. To incorporate reaction distance from equilibrium, we used reversible kinetics called the direct binding modular rate law ^48^ as the enzymatic rate law. Under this framework, a flux *v* can be decomposed into the effect of thermodynamics *η*_rev_ (Fig. 3D) and substrate saturation *η*_sat,s_ ^49^:

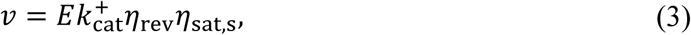

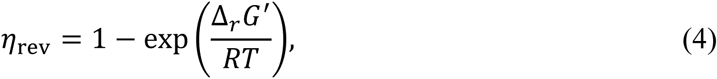

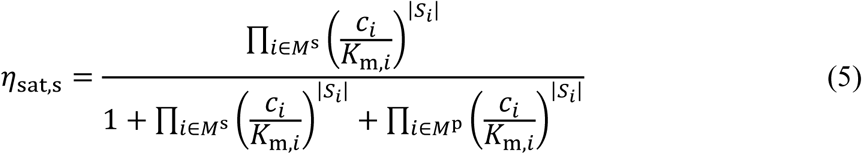

where *E* is an enzyme concentration, *k*_cat_^+^ is the forward catalytic constant, *K*_m_ is the Michaelis-Menten constant, *M*^s^ is substrates, and *M*^p^ is products. Using this reaction rate raw, we calculated FCC based on the connectivity theorem and summation theorem in MCA ^50^ (see Methods). This FCC calculation depends on metabolite concentrations, Δ_*f*_*G*′∘, and *K*_m_. For Δ_*f*_*G*′∘, we used sampling values for the confidence interval calculation in GLEAM. *K*_m_ values were sampled from distributions based on BRENDA database data ^51^. Metabolite concentrations were obtained by the following two approaches.

**Fig. 3:**
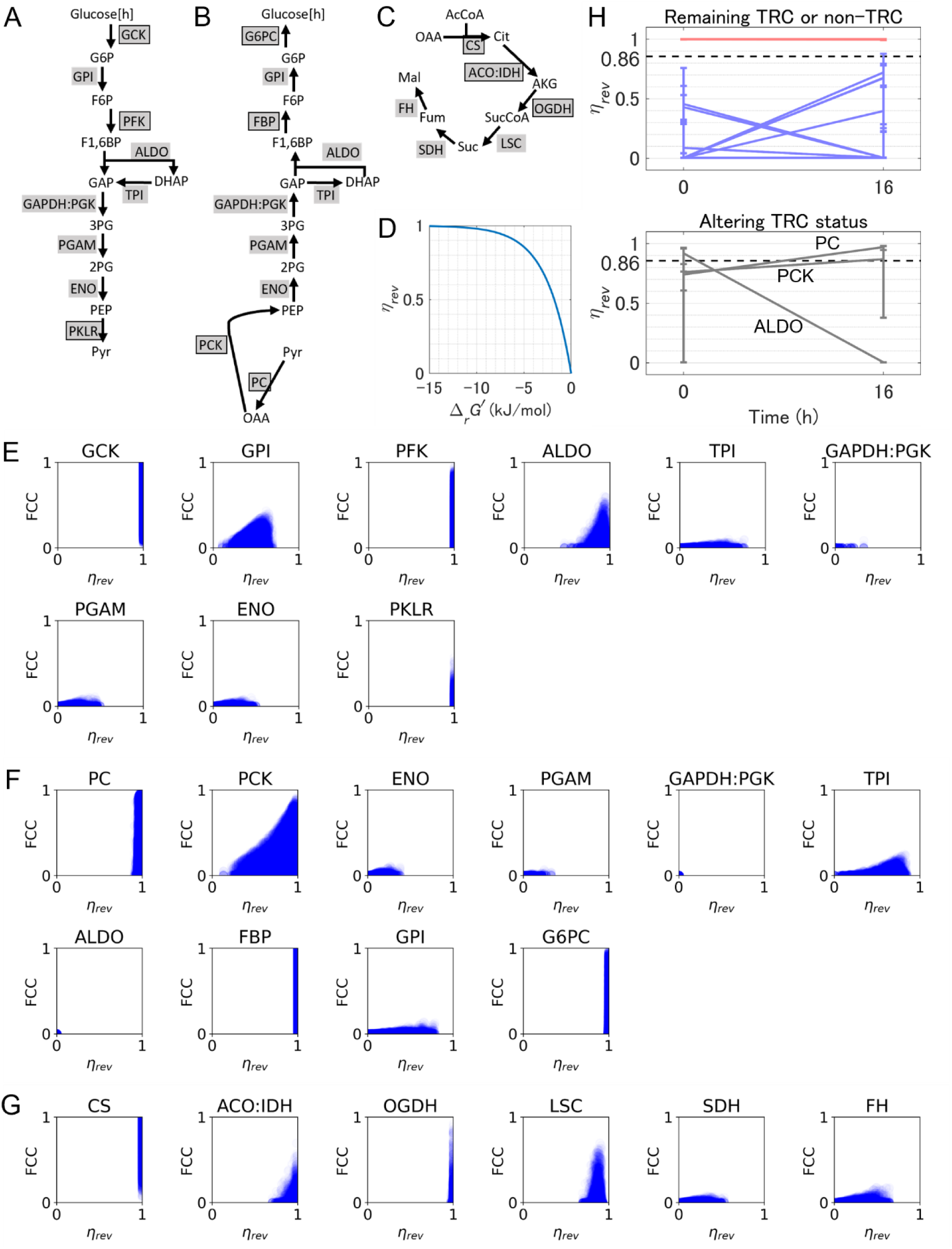
Thermodynamic Rate-limiting Candidates (TRCs) were maintained during fasting. (**A**) Glycolysis pathway where FCCs were calculated. (**B**) Gluconeogenesis pathway where FCCs were calculated. (**C**) The TCA cycle where FCCs were calculated. MDH was excluded from this TCA cycle because it proceeded in the opposite direction to the oxidative TCA cycle. (**D**) Correspondence between Δ_*r*_*G*′ and *η*_rev_. (**E-G**) Relationship between *η*_rev_ and FCC in glycolysis (**E**), gluconeogenesis (**F**), and the TCA cycle (**G**) in the mouse liver. *η*_rev_and FCC values of each point were calculated from each sampled set of metabolite concentrations and Δ_*f*_*G*′∘ values for the calculation of confidence intervals and sampled *K*_m_ values based on BRENDA database (see Methods). Metabolite concentrations at *ad libitum* feeding were used for glycolysis (**E**) and the TCA cycle (**G**), and those after 16-hour fasting were used for gluconeogenesis (**F**). Vertical variation is attributed to differences in *K*_m_ values and condition of the other reactions, while horizontal variation reflects differences in metabolite concentrations and Δ_*f*_*G*′∘. (**H**) Alterations of TRC status based on transitions in *η*_rev_ for reactions in glycolysis and gluconeogenesis before and after starvation. Red, blue, and grey lines depict reactions remaining TRC (5 reactions), reactions remaining non-TRC (10 reactions), and reactions altering their TRC status (either TRC or not) during fasting (3 reactions), respectively. Black dashed lines indicate the boundary between TRC and non-TRC where *η*_*rev*_ = 0.86 *i.e.*, Δ_*r*_*G*′ = −5 kJ/mol. Error bars show 95% confidence intervals.

We investigated FCC for metabolite concentrations in mouse liver using sampled values for confidence interval calculations in GLEAM, and FCC for arbitrary Δ_*r*_*G*′ landscapes to identify thermodynamic conditions for large FCC. FCCs for arbitrary Δ_*r*_*G*′ landscapes showed that FCC value larger than 0.5, which is greater than the sum of FCCs of the other reactions in the pathway, required *η*_rev_ ≈ 1 except for in reactions located upstream in a pathway (Fig. S1 A to D). In mouse liver, reactions with *η*_rev_ ≈ 1 at *ad libitum* feeding included irreversible reactions GCK, PFK, PKLR, and the reversible reaction ALDO in glycolysis. Among them, reactions that could achieve FCC ≈ 1 were GCK and PFK, while the other reactions could achieve FCC of between 0.5 and 0.7 (Fig. 3E). We also found that GPI, which is a reversible reaction located upstream in the pathway, could achieve FCC = 0.42 at *η*_rev_ ≈ 0.70. In gluconeogenesis, reactions with *η*_rev_ ≈ 1 after 16-hour fasting were the irreversible reactions PC, PCK, FBP, and G6PC, all of which could achieve FCC ≈ 1 (Fig. 3F). All the other reactions exhibited only small FCC values. In the TCA cycle, similar FCC distributions were observed at *ad libitum* feeding and after fasting (Fig. 3G, S3). Reactions with *η*_rev_ ≈ 1 were irreversible reactions CS, ACO:IDH, OGDH, and a reversible reaction LSC. Among these reactions, CS and OGDH could achieve FCC ≈ 1, while ACO:IDH and LSC could achieve FCC slightly below 0.7. Generally, reversible reactions have not been considered rate-limiting reactions because they have been recognized to be qualitatively closer to equilibrium than irreversible reactions. However, our quantitative analysis demonstrated that the reversible reactions GPI, ALDO, and LSC could achieve FCC of between 0.4 and 0.7 comparable to several irreversible reactions. Considering the results of arbitrary Δ_*r*_*G*′ landscapes, these large FCC values of the reversible reactions can be attributed to GPI being located relatively upstream in glycolysis, and ALDO and LSC being sufficiently away from equilibrium to exhibit *η*_rev_ ≈ 1 despite being closer to equilibrium than irreversible reactions.

As described above, a reaction with FCC larger than 0.5 seems to be either away enough from equilibrium to exhibit *η*_rev_ ≈ 1, or located upstream in a pathway. Because the location of a reaction in a pathway depends on the definition of the pathway, a reaction being away enough from equilibrium to exhibit *η*_rev_ ≈ 1 is a key determinant for the reaction to have large FCC, sometimes being referred to as the rate-limiting step. Therefore, we introduced Thermodynamic Rate-limiting Candidate (TRC) as reactions with *η*_rev_ ≥ 0.86 corresponding to Δ_*r*_*G*′ ≤ −5 kJ/mol, and examined which reaction is thermodynamically capable of being a rate-limiting step for any pathway definitions at *ad libitum* feeding (Fig. 3H, 0h) and after fasting (Fig. 3H, 16h). TRC reactions both at *ad libitum* feeding and after fasting included GCK, PFK, CS, and seven other reactions, indicating these reactions can be a rate-limiting step regardless of fasting levels. The parameter *η*_rev_ and TRC status (either TRC or not) are sensitive to Δ_*r*_*G*′ variations at a near-equilibrium region (Δ_*r*_*G*′ > −5 kJ/mol) while they are insensitive at a far-from-equilibrium region (Δ_*r*_*G*′ < −10 kJ/mol) (Fig. 3D). Due to this property, 15 out of 18 reactions remained either TRC (red line in Fig. 3H) or non-TRC (blue line in Fig. 3H) across at *ad libitum* feeding and after fasting in glycolysis and gluconeogenesis, where only Δ_*r*_*G*′ of near-equilibrium reactions were strictly maintained. This result suggests that fasting-induced fluctuations in metabolite concentrations occur within a range that maintains the TRC status for most reactions. The three reactions of ALDO, PC, and PCK were the exceptions that did not remain TRC or non-TRC (gray line in Fig. 3H), and whether these reactions are rate-limiting or not would depend on fasting levels. The reactions maintaining their TRC status described above remain stable across the threshold from *η*_rev_ = 0.73 to 0.99.

### Glycolysis and gluconeogenesis achieve efficient directional switching by expending large Enzyme Costs

A fundamental question in biology is to understand what biological systems are optimized for. While objective functions optimized in microbial metabolism—such as maximizing growth or ATP production ^52^ and minimizing enzyme demands ^42^—have been reported, the objective function for *in vivo* mammalian metabolism remains completely unknown. Therefore, we examined the optimality of metabolism in mouse liver from the aspect of metabolite concentrations and Δ_*r*_*G*′ landscape obtained from GLEAM.

By solving the equation of the direct binding modular rate law (Eq. 3) with respect to the enzyme amount, we obtain

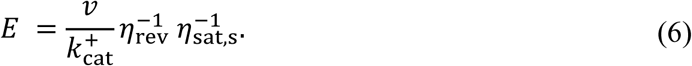

This equation shows that the closer to a reaction is to equilibrium (the smaller *η*_rev_ is), the greater the enzyme amount required to achieve the same flux *v*. Here we defined Enzyme Cost (EC) as the product of enzyme amount and enzyme molecular weight, and Enzyme Cost Minimum (ECM) as the theoretical state that minimizes the total EC within the entire pathway (see Method). By comparing ECs and metabolite concentrations in mouse liver to those in ECM, we investigated whether mouse liver metabolism is optimized for enzyme demands reduction. As with FCC calculation, we calculated EC using sampling values of metabolite concentrations and Δ_*f*_*G*′∘ used in the confidence interval calculation in GLEAM. For *K*_m_ values, we obtained them as geometric means of data in the BRENDA database ^51^.

We calculated EC and ECM for the reactions in glycolysis at *ad libitum* feeding, gluconeogenesis after 16-hour fasting, and the TCA cycle under both these conditions (Table S5). The liver showed larger total EC compared to ECM, with a difference between pathways. The liver/ECM ratio of total EC was on the order of 10^4^ for glycolysis and gluconeogenesis, but only 10^1^ for the TCA cycle (Fig. 4A). Additionally, we compared ECM-predicted metabolite concentrations with liver metabolite concentrations in each pathway, and found typical fold errors of 10^RMSE^ = 29.79 (RMSE: root mean square error on a log10-scale) for glycolysis and gluconeogenesis and 14.40 for the TCA cycle (Fig. 4B). These findings show that enzyme and metabolite concentrations in the liver are not optimized for EC, and this is more evident in glycolysis and gluconeogenesis than in the TCA cycle.

**Fig. 4:**
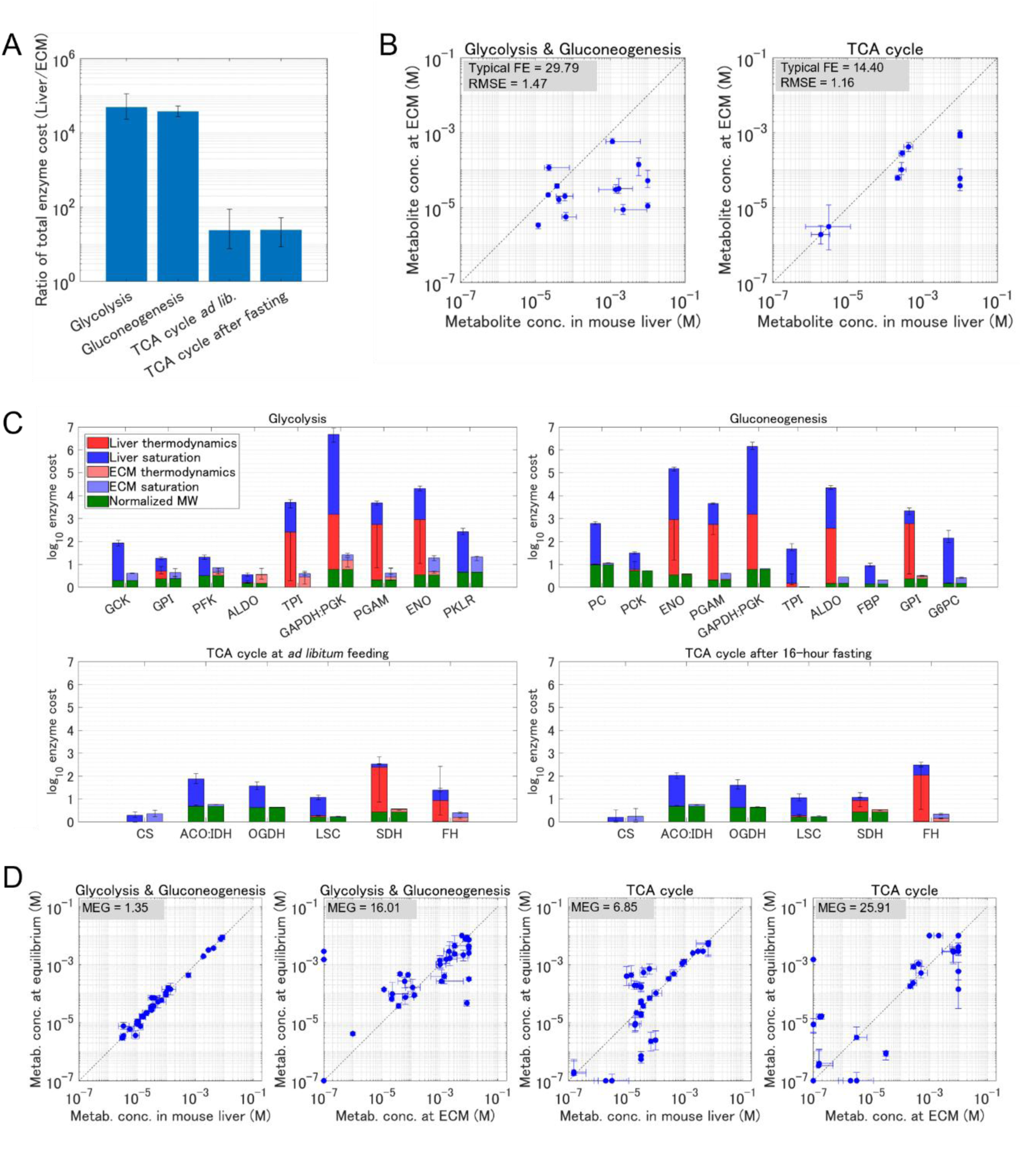
Glycolysis and gluconeogenesis achieve efficient switching by expending large Enzyme Costs. (**A**) Ratio of total enzyme cost under metabolite concentrations in the mouse liver and ECM (Liver/ECM). The total EC was calculated for glycolysis, gluconeogenesis, and the TCA cycle, which are the same as the pathways used for FCC calculation (Fig. 3A to C). Metabolite concentrations at *ad libitum* feeding and after 16-hour fasting were used for glycolysis and gluconeogenesis, respectively. *K*_m_ values for EC calculation were the geometric mean of data in the BRENDA database (Table S5). Error bars show 95% confidence intervals. EC in mouse liver, and metabolite concentrations and EC at ECM are shown in Table S5. (**B**) Comparison of metabolite concentrations in the mouse liver and ECM. Plotted data are for the measured metabolites, and the scatter plot for the TCA cycle includes the data both at *ad libitum* feeding and after 16-hour fasting. RMSE is the root mean square error on a log10-scale of ECM’s prediction compared to metabolite concentrations in the mouse liver, and typical fold error (typical FE) = 10^RMSE^. Error bars show 95% confidence intervals. Plotted data are for the measured metabolites. (**C**) EC of each reaction in glycolysis, gluconeogenesis, and the TCA cycle. The left and right bars of each reaction are EC in the mouse liver and ECM, respectively. Since the vertical axis represents a logarithmic scale, EC can be depicted by the stacked bar of normalized enzyme molecular weight (green), the effect of thermodynamics, *η*_rev_^−1^ (red), and saturation, *η*_sat,s_^−1^ (blue). Note that although the total Gibbs free energy change across the entire pathway in ECM is fixed to that in the mouse liver (see Methods), the sum of *η*_rev_^−1^ differs due to the non-linear increase of *η*_rev_ over Δ_r_*G*′ (Fig. 3D). Error bars show 95% confidence intervals. (**D**) Comparison of metabolite concentrations in the liver or ECM, and those for MEG calculation, where the reactions required to reverse for metabolic switching are at equilibrium. Plotted data are for the measured metabolites. Error bars show 95% confidence intervals. Metabolite concentrations where the reactions required to reverse for metabolic switching are at equilibrium are shown in Table S6.

To identify a key reaction contributing to the large total EC of glycolysis and gluconeogenesis in mouse liver, we broke down the total EC of pathways into ECs of individual reactions (Table S5). The GAPDH:PGK reaction in mouse liver exhibited the highest EC of 10^6^ both in glycolysis and gluconeogenesis (Fig. 4C), while EC of reactions in the TCA cycle (*e.g.*, SDH, ACO:IDH, and FH) are at most on the order of 10^2^. The EC values of GAPDH:PGK are 99% and 89% of the total EC of glycolysis and gluconeogenesis, respectively, suggesting that GAPDH:PGK outstandingly elevates the total EC of glycolysis and gluconeogenesis. The high EC for GAPDH:PGK both in glycolysis and gluconeogenesis is attributed to two factors. One factor is low substrate saturation (Fig. 4C, blue), which is given by the lower substrate metabolite concentrations (3.4×10^−3^ mM of GAP *ad libitum* feeding and 1.3×10^−2^ mM of 3PG after fasting) than their respective *K*_m_ values (1.9×10^−1^ mM for GAP and 3.5×10^−1^ mM for 3PG), resulting in large *η*^−1^_sat,s_. From Eq. 6, a large *η*^−1^_sat,s_, indicating low substrate saturation, causes an increase in EC to carry a certain flux. Under this condition, an increase in flux from higher enzyme levels is mitigated by fluctuations in metabolite concentrations. The other factor of the high EC for GAPDH:PGK is liver thermodynamics (Fig. 4C, red), where the reaction was markedly close to equilibrium (Δ_*r*_*G*′ = −0.01 kJ/mol both at *ad libitum* feeding and after fasting), leading to large *η*^−1^. When the reaction is close to equilibrium, increasing the enzyme abundance enhances both the forward and reverse fluxes, thus limiting the net flux increase. Thus, the elevated EC for this reaction is a result of both low substrate saturation and high reaction reversibility.

The liver glycolysis and gluconeogenesis required much larger EC than ECM, especially in the midstream reversible reactions due to large *η*^−1^ values. We hypothesized that the reason for this may be that the enzyme costs are required for enabling switching from glycolysis to gluconeogenesis during fasting. To assess the metabolic hurdle for directional switching from glycolysis to gluconeogenesis, we introduced the Metabolite Equilibrium Gap (MEG) as the typical fold change in metabolite concentration required for the directional switching (Table S6) (see Methods). In glycolysis and gluconeogenesis, mouse liver exhibited MEG = 1.35 while ECM exhibited MEG = 16.01 (Fig. 4D), suggesting that the liver achieves directional switching from glycolysis to gluconeogenesis with much smaller metabolite concentration changes than ECM. Unlike glycolysis and gluconeogenesis, the TCA cycle, which does not undergo directional switching during fasting, exhibited a larger MEG (6.85) in the liver than that of glycolysis and gluconeogenesis. The mouse liver expends a much larger enzyme cost in glycolysis and gluconeogenesis than the theoretical minimum, allowing it to achieve directional switching with smaller changes in metabolite concentration. Conversely, a smaller enzyme cost is expended for the TCA cycle—which does not undergo directional switching—than for glycolysis and gluconeogenesis.

### Robustness of **Δ_*r*_*G*′** in the liver against obesity

Obesity disrupts liver metabolism, leading to impaired homeostasis in response to fasting ^7,8,10^ and subsequent development of various diseases ^9,18–20^. Recent studies have reported that the obesity-associated dysregulation of liver metabolism occurs even at *ad libitum* feeding ^11,12^. To reveal the impact of obesity-induced metabolic disruption on the thermodynamic characteristics of reactions, we calculated Δ_*r*_*G*′ in the *ob*/*ob* mouse liver during fasting.

The Δ_*r*_*G*′ landscape in *ob*/*ob* mouse liver (Fig. 5A; Table S2) showed that the difference in Δ_*r*_*G*′ from WT mouse liver was less than 5 kJ/mol for most reactions both at *ad libitum* feeding and after 16-hour fasting (Fig. 5B, C, S3). Consequently, the overall trend of irreversible reactions being far from equilibrium (Δ_*r*_*G*′ < −10 kJ/mol) and reversible reactions being near equilibrium (Δ_*r*_*G*′ > −5 kJ/mol) was preserved even under obesity. The similar Δ_*r*_*G*′ values between WT and *ob*/*ob* mouse indicate that the concentration ratios of substrates and products for the reactions are comparable between the genotypes. Individual metabolite concentrations (Table S2), however, showed discrepancies between WT and *ob*/*ob* mouse liver (Fig. 5D), with log2 fold change exceeding 0.5 for 16/31 and 13/31 metabolite concentrations at *ad libitum* feeding and after 16-hour fasting, respectively. This result suggests that the thermodynamic characteristics of reactions remain robust against obesity despite the changes in individual concentrations. Particularly, in most reversible reactions of glycolysis and gluconeogenesis, the differences in Δ_*f*_*G*′ between WT and *ob*/*ob* mouse were not reflected in the differences in Δ_*r*_*G*′ (Fig. 5E). This result prominently demonstrates that the reaction thermodynamic characteristics are robust to obesity while the individual metabolite concentrations are sensitive.

**Fig. 5:**
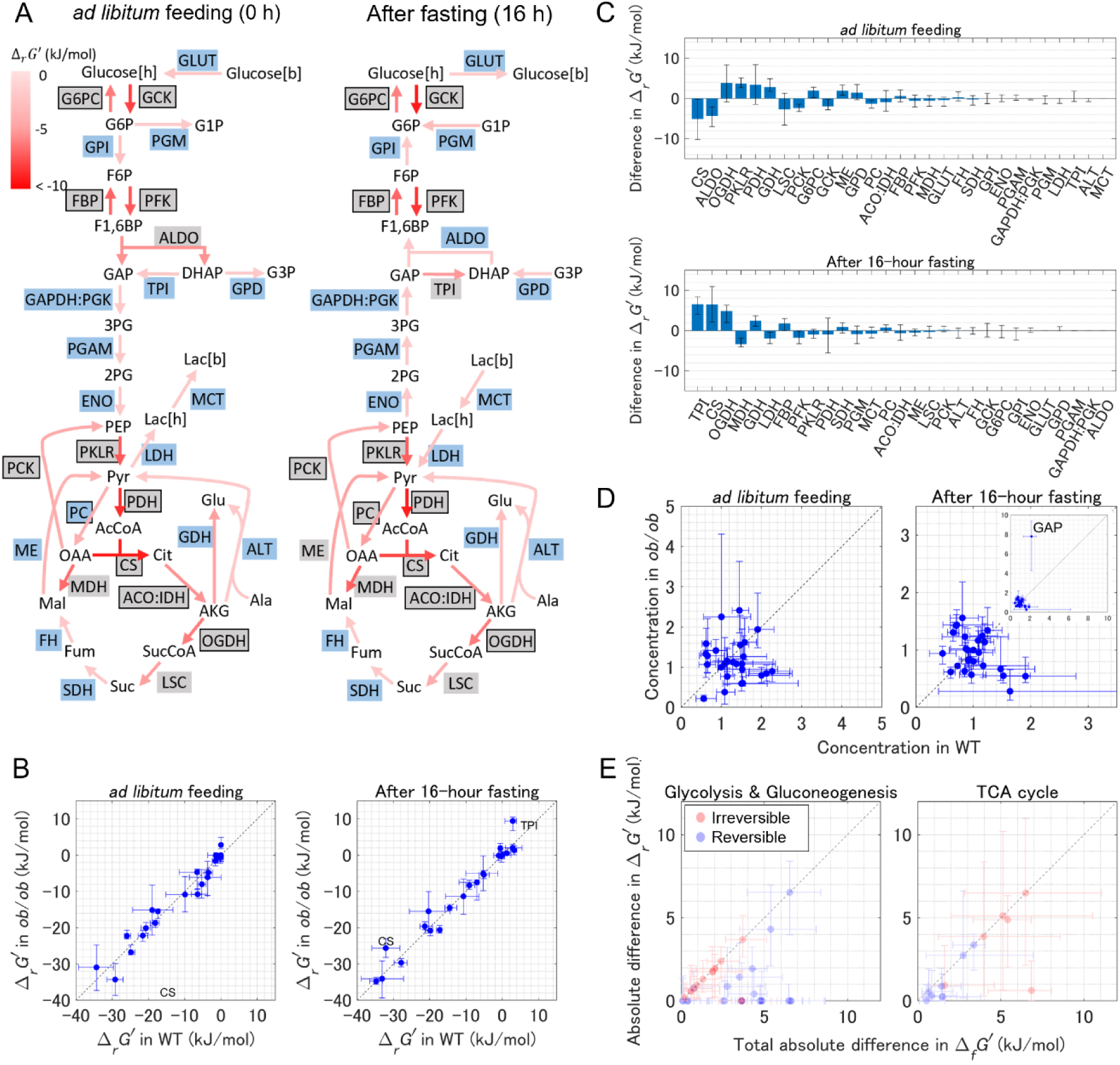
Robustness of Δ_*r*_*G*′ in the liver against obesity. (**A**) Δ_*r*_*G*′ landscape of glucose metabolism in *ob*/*ob* mouse liver at *ad libitum* feeding and after 16-hour fasting. Δ_*r*_*G*′ is depicted by arrow colors. Shaded texts represent reaction names; boxed reactions are irreversible, while unboxed reactions are reversible. Blue shadings indicate near-equilibrium reactions with Δ_*r*_*G*′ > −5 kJ/mol, and gray shadings indicate the other reactions with Δ_*r*_*G*′ ≤ −5 kJ/mol. Unshaded texts represent metabolites. The estimated metabolite concentrations, Δ_*f*_*G*′∘, and Δ_*r*_*G*′ values by GLEAM are shown in Table S2. (**B**) Comparison of Δ_*r*_*G*′ in WT and *ob*/*ob* mouse liver. The forward direction (Δ_*r*_*G*′ < 0) is defined as the reaction direction in WT at *ad libitum* feeding. Error bars show 95% confidence intervals. (**C**) Difference in Δ_*r*_*G*′ between WT and *ob*/*ob* mouse liver (Δ_*r*_*G*′_*ob*/*ob*_ − Δ_*r*_*G*′_WT_). Reactions are arranged from left to right in order of decreasing absolute difference in Δ_*r*_*G*′. Error bars show 95% confidence intervals. (**D**) Comparison of metabolite concentrations in WT and *ob*/*ob* mouse liver. Plotted data represent the concentrations normalized by the mean value at all seven time points and two genotypes *(i.e.*, 14 conditions) to eliminate scale differences in concentrations among metabolites. Error bars show 95% confidence intervals. (**E**) Comparison of absolute difference in Δ_*r*_*G*′ variation and total absolute deference in Δ_*f*_*G*′ = ∑|*S*(Δ_*f*_*G*′_*ob*/*ob*_ − Δ_*f*_*G*′_WT_)|, between WT and *ob*/*ob* mouse liver. Plotted data includes those at *ad libitum* feeding and after 16-hour fasting. The reactions for glycolysis and gluconeogenesis, and the TCA cycles are the same as Fig. 2B. Error bars show 95% confidence intervals.

## DISCUSSION

We revealed the Δ_*r*_*G*′ landscape of glucose metabolism in *in vivo* mouse liver for the first time from fasting time-series absolute quantitative metabolome data using the newly developed GLEAM. In mouse liver, overall trend of Δ_*r*_*G*′ landscape was maintained regardless of fasting level (Fig. 1B, C). The Δ_*r*_*G*′ maintenance was markedly strict in near-equilibrium reactions in glycolysis and gluconeogenesis such as GAPDH:PGK (Fig. 2B), despite large fluctuations in metabolite concentrations (Fig. 1D, 2C), while this strictness was not true for the other reactions such as GCK. In the TCA cycle, overall Δ_*r*_*G*′ variations were small due to minimal metabolite concentration fluctuations during fasting (Fig. 2B, C). This thermodynamic robustness against fasting led to maintenance of the reactions that could have large FCC *i.e.*, TRC during fasting (Fig. 3H) because FCC is sensitive to Δ_*r*_*G*′ variation only in the near-equilibrium region. The thermodynamic landscape in the liver could also contribute to an efficient switching from glycolysis to gluconeogenesis by expending a larger enzyme expression than that theoretically minimized (Fig. 4A to D). We also found similar Δ_*r*_*G*′ landscapes between WT and *ob*/*ob* mouse (Fig. 5A to C), while metabolite concentrations were quite different (Fig. 5D). Our results revealed that the liver robustly maintains Δ_*r*_*G*′ landscapes favorable for metabolic control in contrast to fluctuations in individual metabolite concentrations caused by fasting and obesity.

Methods for estimating thermodynamically consistent metabolite concentrations, Δ_*f*_*G*′∘, and Δ_*r*_*G*′ from absolute quantitative metabolome data have been proposed in previous studies ^26,27,53^. In these methods, Δ_*f*_*G*′∘ values were fixed to the inputs of the model, or estimated without considering the covariance. However, metabolites sharing common chemical groups, such as ADP and ATP, tend to exhibit similar Δ_*f*_*G*′∘ values, because Δ_*f*_*G*′∘ depends on structures of metabolites. Incorporating covariance information into the estimation thus better reflects the underlying chemical relationships between metabolites and allows for narrower confidence intervals in the estimated values ^54^. By leveraging the covariance matrix of Δ_*f*_*G*′∘ from eQuilibrator ^55^, GLEAM achieves more precise and consistent estimation of thermodynamic characteristics throughout the entire metabolic network.

Our study narrowed down the reactions that could achieve large FCC in the pathways, *i.e.*, candidates for a rete-limiting step using the obtained Δ_*r*_*G*′ landscape in mouse liver (Fig. 3E to G, S2). A previous study experimentally measured FCC in glycolysis in iBMK cells, and reactions exhibiting large FCCs are hexokinase (HK), which catalyzes the same reaction as GCK, and PFK ^56^. Another previous study exhibited PC having dominant FCC in gluconeogenesis in rat liver ^57,58^. Although no studies have measured FCC in the TCA cycle to our knowledge, CS, IDH, and OGDH have conventionally been regarded as rate-limiting steps. All these reactions are included in the TRCs in this study, supporting the validity of our Δ_*r*_*G*′ estimation even though this study focuses on mouse liver metabolism. While one study based on standard-state calculation has indirectly showed the connection between reactions away from equilibrium and driver nodes to control the entire network defined using control theory in terms of topological features ^59^, the functional advantage of controlling pathway flux via these reactions remains unclear and is an important future topic for understanding system-level metabolic regulation.

When calculating EC, we made two assumptions: having common turnover number, *k*_cat_ values for all reactions and setting saturation (*c*/*K*_m_) to the geometric mean of the pathway when *K*_m_ could not be obtained from BRENDA. These assumptions were required because of the difficulty of obtaining kinetic parameters in mammals. The former assumption may cause inaccurate weighting of individual enzymes in the total EC minimization, and the latter may lead to inaccurate *η*^−1^_sat,s_. However, the metabolite concentrations close to ECM were reported with a modeling where *k*_cat_ and *K*_m_ information was lacking and only the influence of *η*_rev_^−1^ was considered ^42^. This means that the Δ_*r*_ *G*′ landscape in ECM would be similar under any *k*_cat_ and *K*_m_ values. Therefore, our conclusion of mouse liver metabolism not being optimized for enzyme cost reduction would be reasonable regardless of *k*_cat_ and *K*_m_ assumptions.

According to Eq. 23, 24, 30 to 35, and 38, the design of Δ_*r*_*G*′ landscape faces a trade-off between FCC concentration and enzyme cost minimization. If only a few reactions in a metabolic pathway are far from equilibrium and exhibit *η*_rev_ close to 1, large FCCs are likely to be concentrated in these reactions. In this case, *η*_rev_ of other reactions are close to zero, leading to a large EC. This trade-off can be rephrased as a trade-off between flexibility responding to multiple conditions and survival cost reduction under a single condition because the pathway flux can be efficiently regulated through a minimal number of enzyme activity changes when FCCs are concentrated in limited reactions. A similar trade-off also occurs between MEG reduction and enzyme cost minimization. In mouse liver, particularly in glycolysis and gluconeogenesis, reactions that could achieve high FCC were concentrated in a few reactions (Fig. 3E to G, S2), and MEG was remarkably small (Fig. 4D), while there were large deviations between mouse liver and ECM in total EC and metabolite concentrations (Fig. 4A, B). These results indicate that metabolic regulation in mouse liver adopts a strategy of flexibly responding to changes in environmental and physiological conditions rather than optimizing for a certain condition. In contrast, the previous study in *E. coli* showed that both enzyme concentrations and metabolite concentrations are well predicted by ECM within a typical fold error range of 10^0 42^ (Fig. S3). Taken together, these facts suggest differences in metabolic regulation strategies between cell types; mouse liver expends costs to perform flexible metabolic regulation, while *E. coli* optimized survival cost in a certain condition. Furthermore, our study also showed that this difference in metabolic regulation strategy may also exist between pathways by demonstrating the large differences in TRC proportions and total EC optimality between glycolysis and gluconeogenesis, and the TCA cycle. Note that although glycolysis in mouse liver and *E. coli* shared a common thermodynamic tendency of reversible reactions being close to equilibrium with Δ_*r*_*G*′ < −10 kJ/mol (Fig. 1E), EC in them can be greatly different. As shown in Eq. 38 and 39, thermoduric characteristics of reactions affect EC through *η*^−1^, which indicates *v*^total^/*v*^net^, where *v*^total^ is the total flux of forward and backward reactions, and *v*^tatal^ is the net flux. In *E. coli*, most reversible reactions exhibited Δ_*r*_*G*′ < −1 kJ/mol, contributing to reducing EC with *η*^−1^ on the order of 10^0^ (Fig. S4, orange) ^26,60^. In contrast, most reversible reactions in mouse liver exhibited Δ_*r*_*G*′ > −0.1 kJ/mol, where *η*^−1^ rises sharply (Fig. S4, green), leading to much larger EC than *E. coli*.

One of the limitations of this study is that we assumed hepatocytes as a single compartment and calculated Δ_*r*_*G*′ from metabolite concentrations averaged within the cell. As glycolysis operates in the cytosol and the TCA cycle operates in mitochondria, more accurate Δ_*r*_*G*′calculation requires concentrations in each of these compartments. Furthermore, localized concentrations can be affected by substrate channeling ^61^. Substrate channeling is a phenomenon where enzymes of consecutive reactions form complexes, allowing the product of the former enzyme to be directly transferred to the active site of the latter enzyme without being released into the solvent. For reactions involved in substrate channeling, Δ_*r*_*G*′ is dependent on the channeled metabolite concentration in a limited space among the enzymes. Although substrate channeling has not yet been confirmed in living organisms, the channeling of oxaloacetate between MDH and CS has been indirectly supported by experiments ^62,63^, and ALDO, TPI, and GAP have also been suggested to form complexes ^64,65^. Another limitation is that the FCC calculations were based solely on stoichiometric relationships and did not account for allosteric regulations because of the severe lack of *K*_i_ in mammals and simplification of the calculation. Inclusion of allosteric regulations would alter FCC values, allowing for further narrowing down the reactions that can have a large FCC.

In conclusion, we developed GLEAM to estimate Δ_*r*_*G*′ in the glucose metabolism in mouse liver and revealed the thermodynamic robustness against fasting and obesity. Furthermore, we suggested that these thermodynamic characteristics in the liver contribute to efficient metabolic control in two ways: the maintenance of rate-limiting step candidates and the switching between glycolysis and gluconeogenesis with minimal metabolite fluctuations by expending large enzyme expressions. This study thus provides crucial insights into the principles underlying the highly flexible and robust metabolic regulation observed in mammalian liver, paving the way for future investigations into therapeutic targets for metabolic diseases.

## RESOURCE AVAILABILITY LEAD CONTACT

Requests for further information and resources should be directed to and will be fulfilled by the lead contact, Shinya Kuroda.

## MATERIAL AVAILABILITY

This study did not generate new unique reagents.

## DATA AND CODE AVAILABILITY

Metabolomic data from WT and *ob*/*ob* mice in our previous studies ^8,66^ were used in this study.

The information on kinetic parameter (Michaelis-Menten constant and turnover number) distributions used for FCC calculation, the FCC values generated in this study, and the MATLAB code for GLEAM and the analysis of FCC and EC are deposited in GitHub (https://github.com/tabekawa/Thermodynamics_MouseLiver). All other data needed to evaluate the conclusions in the paper are present in the paper or the supplemental information.

Any additional information required to reanalyze the data reported in this paper is available from the lead contact upon request.

## ACKNOWLEDGMENTS

We thank our laboratory members for critically reading this manuscript. The computational analysis of this work was performed in part with support of the supercomputer system of the National Institute of Genetics (NIG), Research Organization of Information and Systems (ROIS).

This study was supported by the Japan Society for the Promotion of Science (JSPS) KAKENHI (Grant Numbers JP17H06300, JP17H06299, JP18H03979, JP21H04759, JP22K17992, JP23H04946, JP23H04939) (S.K.); the Japan Science and Technology Agency (JST) as part of CREST (JPMJCR2123, JPMJCR25T3) and as part of ASPIRE (JPMJAP24B1) (S.K.); The Uehara Memorial Foundation (S.K.); AMED Grant Number JP21zf0127001 (T.S.); JST, CREST Grant Number JPMJCR2123 (T.S.); MEXT KAKENHI Grant Number JP23H04946 (T.S.); World Premier International Research Center Initiative (WPI), Human Biology-Microbiome-Quantum Research Center (Bio2Q), MEXT, Japan (T.S.); JSPS KAKENHI JP24K11417 (A.H.); and the FOREST Program of the Japan Science and Technology Agency (JPMJFR2052) (A.H.).

## AUTHOR CONTRIBUTIONS

Conceptualization: T.A., S.O., and S.K. Methodology: T.A., S.O., A.H., and T.S. Investigation: T.A., A.H., and T.S. Visualization: T.A.

Funding acquisition: A.H., T.S., and S.K. Project administration: S.K. Supervision: S.K.

Writing – original draft: T.A., S.O., and S.K.

Writing – review & editing: T.A., S.O., A.H., T.S., and S.K.

## DECLARATION OF GENERATIVE AI AND AI-ASSISTED TECHNOLOGIES

During the preparation of this work, the authors used ChatGPT, DeepL, and Claude in order improve the readability and language of the manuscript. After using these tools, the authors reviewed and edited the content as needed and take full responsibility for the content of the publication.

## STAR METHODS

### EXPERIMENTAL MODEL AND PARTICIPANT DETAILS

Metabolomic data from WT and *ob*/*ob* mice in our previous studies ^8,66^ were used in this study.

### METHOD DETAILS

#### Metabolic network for glucose metabolism in mice

A metabolic network for glucose metabolism in mice was constructed to calculate Δ_*r*_*G*′. The network consists of total 55 reactions and 80 metabolites in the liver, muscle, and blood (the same reactions and metabolites in different compartments were counted as distinct ones), and involves glycolysis, gluconeogenesis, TCA cycle, glycogen metabolism, alanine metabolism, and transporters (Fig. S5, Table S3). Among the 55 reactions, we defined 23 reversible reactions and 32 irreversible reactions based on the tissue-specific genome-scale metabolic network ^67^. All the transporters were defined as reversible reactions.

Reversible reactions GAPDH and PGK were combined into a single reversible reaction GAPDH:PGK because 1,3-bisphosphoglycerate was not measured in the metabolomic data^8^. According to the experimentally measured metabolite concentrations, a reversible reaction ACO proceeded in the reverse direction of the oxidative TCA cycle with a little thermodynamic driving force at all the time points in WT and *ob*/*ob* mice, breaking a sequential pathway from CS to FH. Since this long sequential pathway is necessary for the downstream analysis, such as FCC and MEC calculation, we summed ACO and irreversible reaction, IDH, which have a large thermodynamic driving force in the direction of the oxidative TCA cycle, as a single irreversible reaction, ACO:IDH.

In the skeletal muscle, FBP, G6PC, and PCK in gluconeogenesis were not included in the metabolic network. Although PC is also involved in gluconeogenesis, PC is included in the metabolic network of skeletal muscle because we could not exclude the possibility of its expression in the skeletal muscle ^68,69^.

In this study, we assumed the liver and skeletal muscle as a single compartment of hepatocytes and myocytes, respectively, and cytosolic and mitochondrial compartments in these cells were not considered for simplification of the modeling. Therefore, the Δ_*r*_*G*′calculated in this study represents the average values within the cells.

#### Algorithm of GLEAM

We developed GLEAM as a method to estimate complete set of thermodynamically consistent metabolite concentrations and Δ_*f*_*G*′∘ values, from metabolomic data on a logarithmic scale, 𝐥𝐧 𝒄 ∈ ℝ^*m*×*h*×*g*^, and Δ_*f*_*G*′∘ data calculated by the component contribution method ^70^, 𝚫_𝒇_𝑮′∘ ∈ ℝ^𝑚^, where *m*, *h*, *g* are the number of metabolites, timepoints, and genotypes, respectively. Since metabolomic and Δ_*f*_*G*′∘ data contain uncertainties and missing values, directly substituting them into the definition equation of Δ_*r*_*G*′ (Eq. 1, 2) often violates the second law of thermodynamics. To address this issue, GLEAM corrects for the uncertainties and missing values by calculate the most plausible values based on measured metabolomic data within a thermodynamically feasible space.

We first formulate the quadratic programming problem to minimize the departures of estimated metabolite concentrations and Δ_*f*_*G*′∘ from the experimental and database values under thermodynamic and physiological constraints:

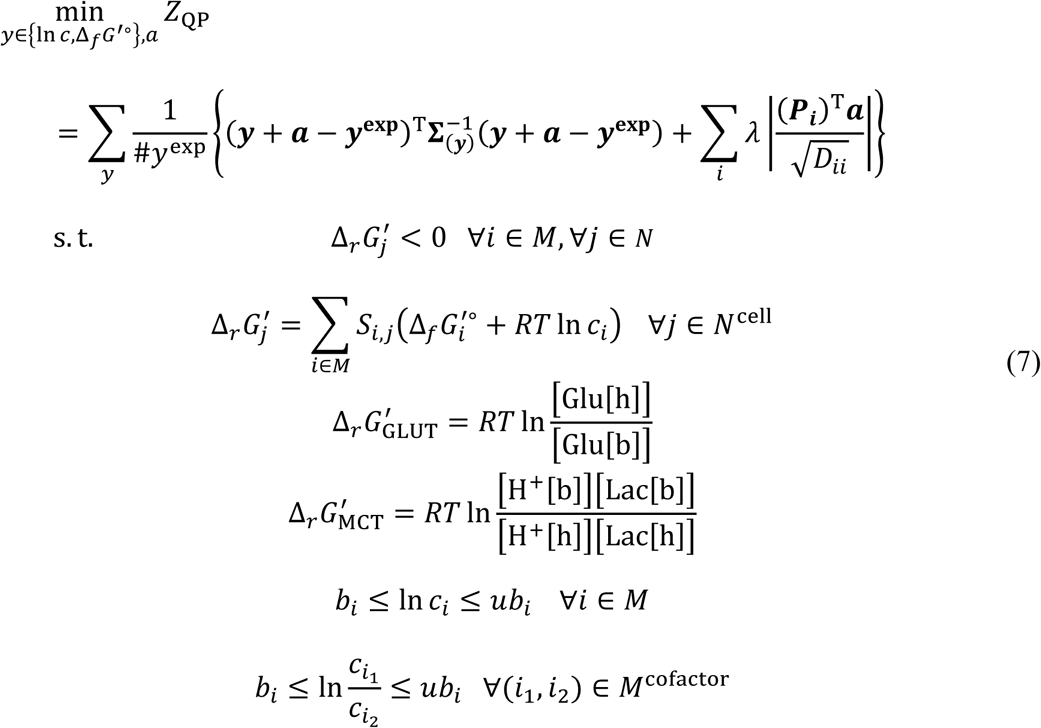

where *c*_*i*_ is concentration of metabolite *i*, *S*_*i*𝑗_ is stoichiometric coefficient of metabolite *i* in reaction *j*, *R* is gas constant, and *T* is temperature. *M*, 𝑁, and 𝑁^cell^ are sets of metabolites, reactions, and intracellular reactions, respectively. *M*^cofactor^ is the cofactor pairs of which the ratio was constrained. 𝚺 is variance-covariance matrix of which the eigen decomposition is 𝚺 = 𝐏𝐃𝐏^T^. 𝑎 is systematic error, and 𝜆 is regularization parameter. The superscript exp means measured or database value. The first term in curly brackets of the objective function represents the variance and covariance-weighted residuals of estimated metabolite concentrations and Δ_*f*_*G*′∘, which considers systematic errors from experimental operations, while the second term represents a regularization term for systematic errors to avoid overfitting. By using the covariances in Δ_*f*_*G*′∘ for the weight of residuals, the estimation reflects the correlations in Δ_*f*_*G*′∘ due to shared chemical groups. In contrast, all covariances between metabolite concentrations were set to zero, making 𝚺 a diagonal matrix, since no prior correlation in metabolite concentrations is expected.

The solution of the optimization problem (7) may not provide unique concentrations, especially for unmeasured metabolites, which can lead to undetermined Δ_*r*_*G*′ of the reactions involving the unmeasured metabolites. To estimate unique Δ_*r*_*G*′, two terms with variance of unmeasured metabolite concentrations were added to the objective function:

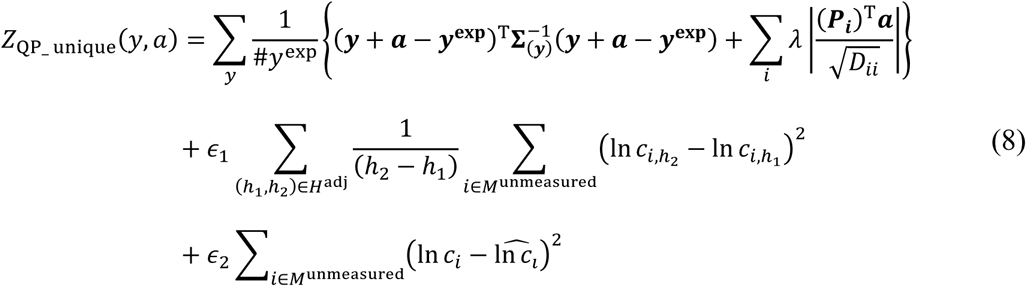

where 𝐻^adj^ is pairs of adjacent timepoints, *M*^unmeasured^ is unmeasured metabolites, 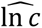 is median of upper and lower bounds of metabolite concentration, and 𝜖_1_ is small values enough to prioritize the first term over the second term. Similarly, 𝜖_2_ (𝜖_1_ ≫ 𝜖_2_) allows the second term to take precedence over the third term. The second term of the objective function (the first additional term) reduces the sum of concentration changes of unmeasured metabolites across time points, making smooth concentration changes more likely to be estimated than unnatural repetition of increases and decreases. The third term of the objective function (the second additional term) departures of the estimated unmeasured metabolite concentrations from the median of their upper and lower bounds. This term can be justified by the trade-off in increasing concentrations between elevation of enzyme efficiency and cellular burden from osmotic pressure and toxicity ^40,71^. Since the objective function includes terms with greatly different orders of magnitude due to 𝜖 ∈ {𝜖_1_, 𝜖_2_}, it has difficulty in optimized at once. Therefore, in implementation, we first solve the optimization problem only for the first term, then, with the measured metabolite concentrations and Δ_*f*_*G*′∘ fixed to the obtained solutions, and then solve the optimization problem for the second and third terms.

Metabolic networks can include reversible reactions and their directions are unknown under a condition of interest. By introducing binary variables representing reaction, GLEAM model is finally formalized by the following mixed integer quadratic programming problem:

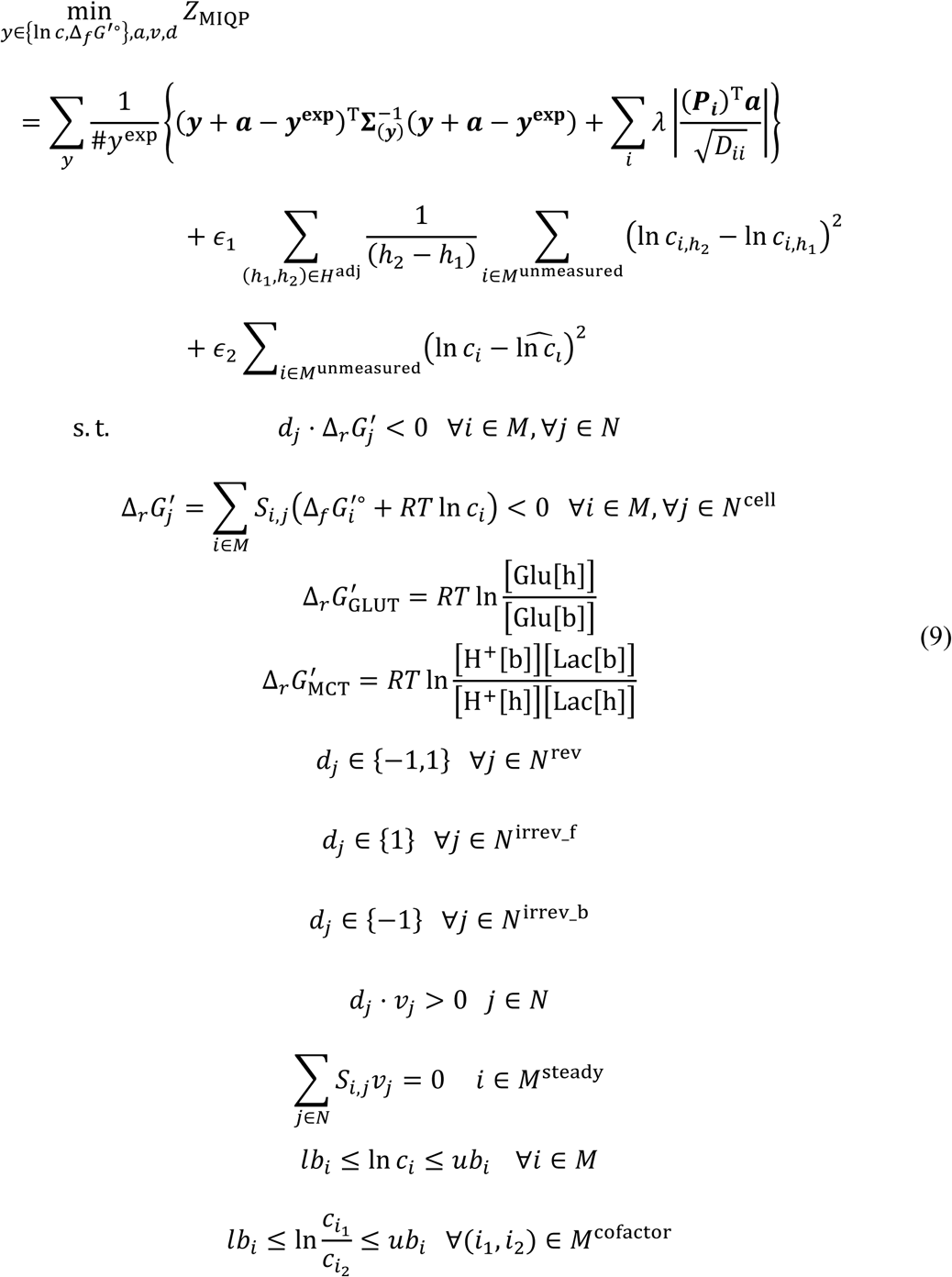

where *v*_𝑗_ is metabolic flux through reaction *j*, 𝑑_𝑗_ = {1, −1} is a binary variable representing direction of reaction *j*, and 𝑁^rev^ is reversible reactions. 𝑁^irrev_f^ and 𝑁^irrev_b^are irreversible reactions of which the direction is fixed to forward and backward with respect to the definition of metabolic network, respectively. *M*^steady^ is metabolites assumed to be in steady state. To solve this mixed integer quadratic programming problem efficiently, GLEAM treats it as a bilevel optimization problem, where the optimization of binary variables is the upper-level problem and the optimization of continuous variables is the lower-level problem:

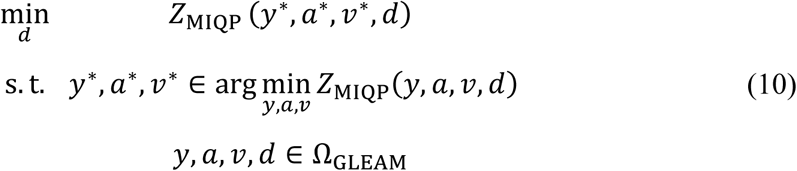

where Ω_GLEAM_ is the solution space defined in the problem (9). In the bilevel optimization problem, binary variables are optimized by a genetic algorithm, while continuous variables are optimized by the interior-point method under the individuals of binary variables (Fig. S6). The genetic algorithm was performed by the MATLAB function ga with termination condition of FunctionTolerance = 10^-9^ and population size of PopulationSize = 500. The lower-level quadratic programming problem was solved using Gurobi ^72^ MATLAB API with dual feasibility tolerance of OptimalityTol = 10^-9^ and primal feasibility tolerance of FeasibilityTol = 10^-9^. This bilevel optimization was performed 10 times from random initial values, and the solution with the minimum objective function value was selected as the optimal solution.

#### Calculation of confidence intervals

The confidence intervals of the estimation by GLEAM were calculated using parametric bootstrap sampling of metabolite concentrations and Δ_*f*_*G*′∘. These data were first resampled by adding perturbation based on the variance and residual of each metabolite concentration and Δ_*f*_*G*′∘ in the Δ_*r*_*G*′ estimation by Eq. 9 using the experimental and database values ^27,73^. To generate Δ_*f*_*G*′∘ samples reflecting their covariance, the perturbations were applied after the basis of Δ_*f*_*G*′∘ vector was transformed by transposed matrix of the eigenvectors of the variance-covariance matrix:

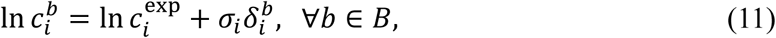

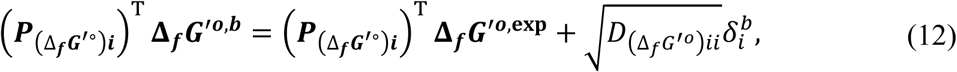

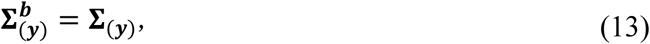

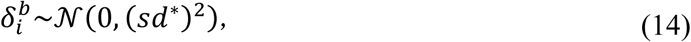

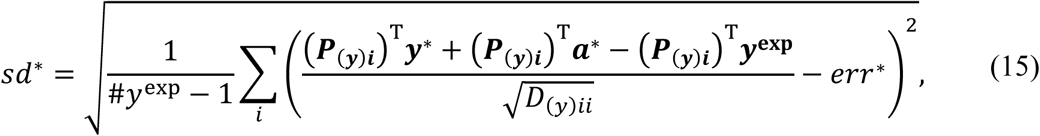

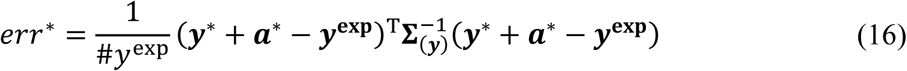

where 𝑏 is the indices of bootstrap samples 𝐵, ∗ represents the optimized values by Eq. 9 using the experimental and database values. 𝛿 is gaussian noise generated from the normal distribution with mean 0 and standard deviation 𝑠𝑑^∗^, which represents the standard deviation of metabolite concentration and Δ_*f*_*G*′∘ residuals in the Δ_*r*_*G*′ estimation from the experimental and database values. Using the resampled metabolite concentrations and Δ_*f*_*G*′∘values, we conducted the estimation of thermodynamically consistent metabolite concentrations and Δ_*f*_*G*′∘ in the same manner as the estimation from the experimental and database values. In this optimization, the directions of reversible reactions were fixed to those estimated from the experimental and database values. This resampling and optimization were performed 2000 times. By applying the percentile method to the resulting 2000 sets of the estimated values, we obtained the 95% confidence intervals.

#### Selection of regularization parameter

The regularization parameter 𝜆 for systematic errors in GLEAM was selected by the reduced chi-square (𝜒^2^_red_). 𝜒^2^_red_ was calculated using the estimated metabolite concentrations and Δ_*f*_*G*′∘ from resampled values ^27,73^:

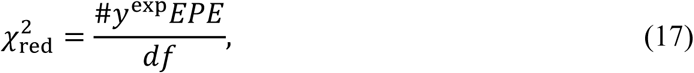

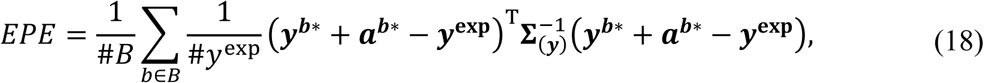

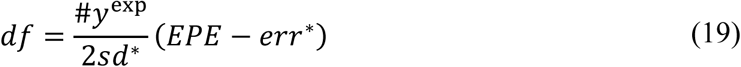

where 𝑏 ∗ represents the optimized values calculated from the resampled values. The model is interpreted as fitting the data ideally when 𝜒^2^_red_ = 1, as overfitting when 𝜒^2^_red_ < 1, and as poorly fitting when 𝜒^2^_red_ > 1. In this study, 𝜒^2^_red_ = 1.03 when 𝜆 = 1.51, and therefore it was selected as the best regularization parameter.

#### Application of GLEAM to mice data

We applied GLEAM to absolute quantitative metabolome data in the liver, skeletal muscle, and plasma from WT and *ob*/*ob* mice (n = 5) after 0, 2, 4, 6, 8, 12, and 16 hours of fasting (Table S1), which were measured in our previous studies ^8,66^. We used the logarithmic mean and variance of the metabolites, which were successfully measured in three or more samples per condition, and metabolites with two or fewer measured samples were treated as unmeasured metabolites. The cofactors ATP, ADP, GTP, GDP, NAD^+^, NADH, FAD, FADH2, NADP^+^, and NADPH were treated as unmeasured metabolites regardless of the number of measured samples due to sample degradation, and instead, their ratios were constrained by previous reports (Table S7). Intracellular lactate was also treated as an unmeasured metabolite in the same manner as the cofactors because it might leach out of the container during measurement. The upper and lower bounds for many metabolites in GLEAM were set to a physiologically typical range of intracellular metabolite concentration (lower bound: 10^-7^ M, upper bound: 10^-2^ M) ^26^, while the bounds for some unmeasured metabolites were individually defined based on biological knowledge (Table S7).

Δ_*f*_*G*′∘ data calculated by the component contribution method ^70^ were retrieved from eQuilibrator database (Table S1) ^55^. We used the identical Δ_*f*_*G*′∘ data for both liver and skeletal muscle under the condition in eQulibrator of pH = 7.2, pMg = 3.0, and ionic strength = 0.15 M ^26^. We also set the condition in blood to pH = 7.4 ^74^ because Δ_*r*_*G*′ for MCT depends on pH in the cells and blood. Temperature was input to *T* = 310.15 K for all the compartments.

In the liver, glycolysis was assumed to be dominant at the 0h-fasting state (*ad libitum* feeding) while gluconeogenesis was assumed to take over after no later than 16h-fasting, both in WT ^75–79^ and *ob*/*ob* mouse ^80^. At these two timepoints, the directions of reversible reactions involved in glycolysis and gluconeogenesis (GPI, ALDO, TPI, GAPDH:PGK, PGAM, ENO, PGM, GPD, LDH) were fixed and treated as those of the irreversible reactions in the calculation of GLEAM.

Since the reaction rates are sufficiently faster than the temporal changes in metabolite and enzyme levels during starvation, our metabolic model was assumed to be in a pseudo-steady state at each measured time point. This assumption leads to mass balance equilibrium for the intermediate metabolites (G6P, F6P, F1,6BP, DHAP, GAP, 3PG, 2PG, PEP, Pyruvate, AcCoA, Cit, Isocit, AKG, SucCoA, Suc, Fum, Mal, OAA) within the metabolic network.

### Derivation of FCC

We derived flux control coefficients (FCCs) within glycolysis (Fig. 3A), gluconeogenesis (Fig. 3B), and the TCA cycle (Fig. 3C) from the direct binding modular rate law ^48^, which is equivalent to the reversible Michaelis-Menten kinetics ^81^ for simple reversible uni-uni reactions, using theorems of Metabolic Control Analysis (MCA) ^36,47^. In the direct binding modular rate law, flux can be described by the effect of the maximal rate, the thermodynamics, and the saturation of substrate ^49^:

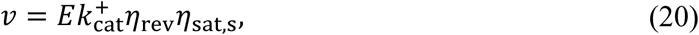

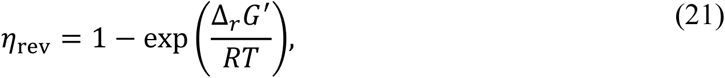

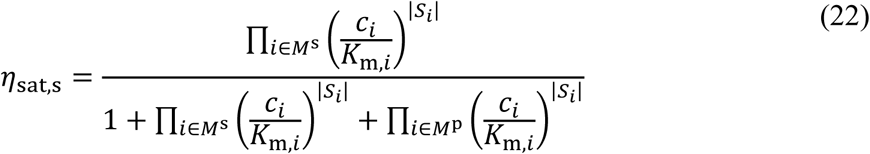

where *M*^s^ is substrates, *M*^p^ is products. From Eq. 20, we calculated the elasticity of substrate 𝜖_s,*i*_^*v*^ and product 𝜖_p,*i*_^*v*^ :

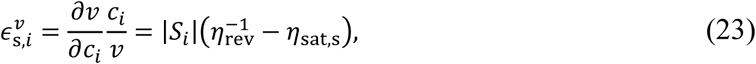

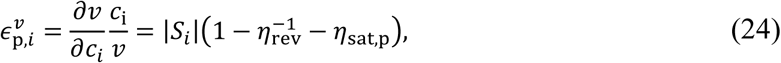

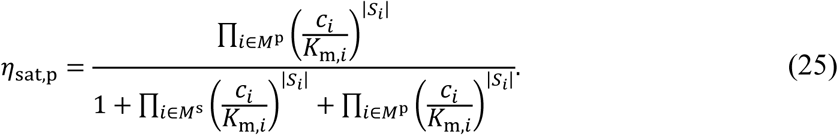

The stoichiometric matrices of the pathways we analyzed here are full rank, and each of their null spaces is one-dimensional, which means the degree of freedom of steady-state flux is one. For these pathways, the following Eq. 26 and 27 hold from the connectivity theorem and the summation theorem in MCA, respectively ^50^:

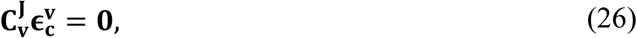

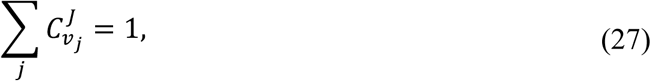

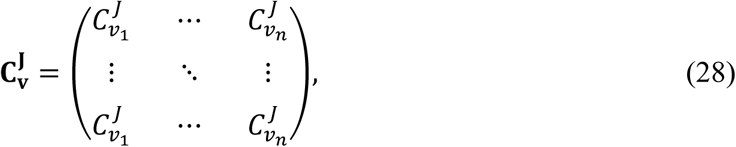

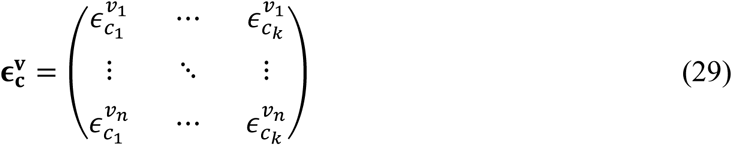

where 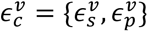. Integrating Eq. 23, 24, 26, and 27, we obtained FCC as:

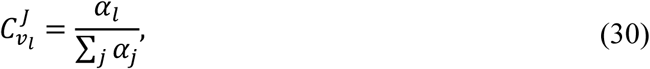

where 𝛼 for GAPDH:PGK in glycolysis is described as

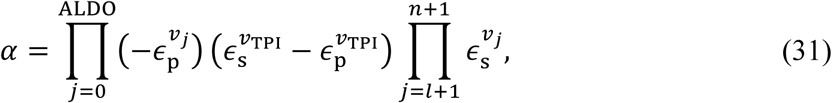

𝛼 for downstream reactions of GAPDH:PGK in glycolysis is described as

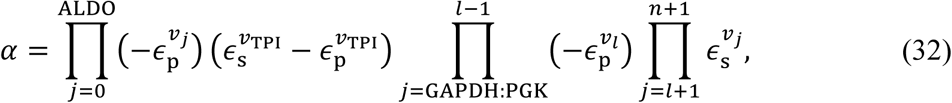

𝛼 for ALDO in gluconeogenesis is described as

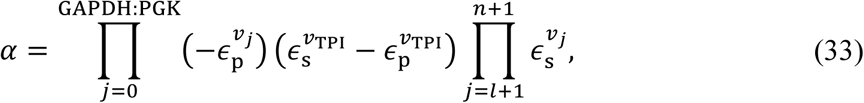

𝛼 for downstream reactions of ALDO in gluconeogenesis is described as

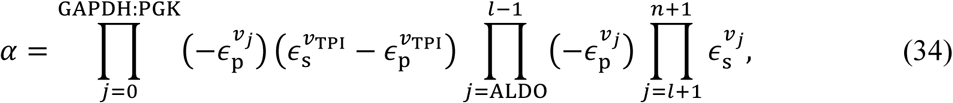

𝛼 for other reactions is described as

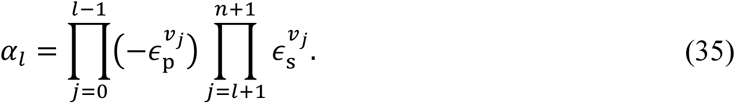

Note that reaction indices 𝑗 ∈ {1, . . ., 𝑁} are denoted such that 1 represents the most upstream reaction and N represents the most downstream reaction in the pathway, and 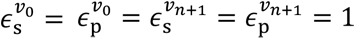.

#### Metabolite concentrations and 𝚫_𝒇_𝑮′∘ values used in FCC calculation

We calculated FCCs under arbitrary Δ_*r*_*G*′ landscapes as well as under the hepatic ones. For both conditions, we used Δ_*f*_*G*′∘ values sampled for confidence interval calculation in GLEAM. To generate the hepatic Δ_*r*_*G*′ landscapes, we used metabolite concentrations sampled in the confidence interval calculation. To generate arbitrary Δ_*r*_*G*′ landscapes, logarithmic metabolite concentrations were uniformly sampled from the physiologically reasonable solution space Ω_ln *c*_:

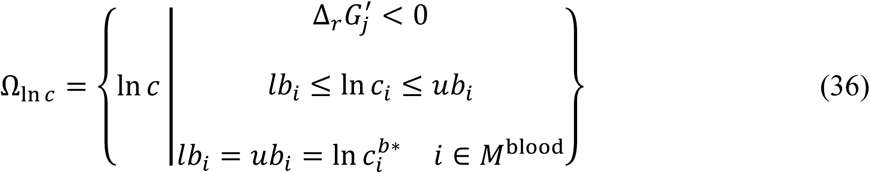

where *M*^blood^ represents blood metabolites. The upper and lower bounds of metabolite concentrations were the same as those used in GLEAM (Table S7). This sampling was performed 20 times for each Δ_*f*_*G*′∘ set using MATLAB function polySampler in the Volume-and-Sampling package ^82^. Thus, we obtained 2000 × 20 = 40000 arbitrary Δ_*r*_*G*′landscapes in total.

#### **𝑲_𝐦_** values used in FCC calculation

In FCC calculation, we sampled Michaelis-Menten constants *K*_m_ for each Δ_*r*_*G*′ landscape to calculate reaction saturation level. To sample *K*_m_, we first retrieved *K*_m_ and the turnover number, 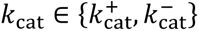 values for enzymes of *Mus musculus*, *Rattus norvegicus*, *Homo sapiens*, *Bos taurus*, *Sus scrofa*, *Canis lupus familiaris*, and *Mammalia* ^83^ from BRENDA database ^51^. For *K*_m_ and *k*_cat_ values from three or more data without reports of mutant enzymes, we calculated their logarithmic means and standard deviations. For *K*_m_ and *k*_cat_from fewer than three data, we set the logarithmic mean of *k*_cat_ (/s) to 1 with a standard deviation of 1.5, and the logarithmic mean of *K*_m_ (mM) to −1 with a standard deviation of 1 ^84^. With the means and standard deviations of *K*_m_ and *k*_cat_, we uniformly sampled logarithmic *K*_m_, *k*_cat_ values for each enzyme from the solution space Ω_KP_:

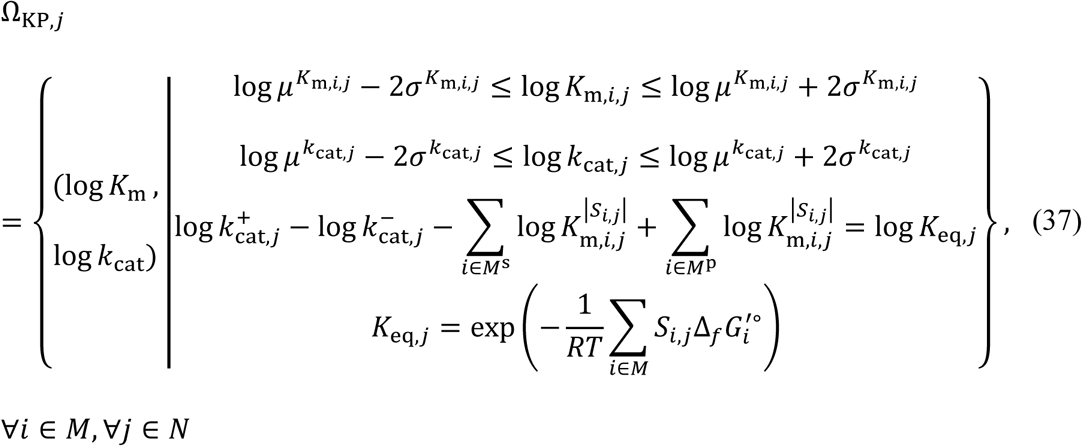

where 𝜇 is mean, 𝜎 is standard deviation, and *K*_eq_ is equilibrium constant. The third constraint reflects the Haldane relationship ^81^. These samplings were performed 20 times for each Δ_*r*_*G*′ landscape using MATLAB function polySampler in the Volume-and-Sampling package ^82^. Thus, we finally calculated FCC under the arbitrary and hepatic Δ_*r*_*G*′ landscapes with 20 × 20 × 2000 = 800000 and 20 × 2000 = 40000 sets of metabolite concentrations, Δ_*f*_*G*′∘ and *K*_m_ values, respectively.

#### Definitions of EC and total EC

By solving the equation of the direct binding modular rate law (Eq. 20) with respect to enzyme amount, we can obtain enzyme demand needed to carry a certain flux:

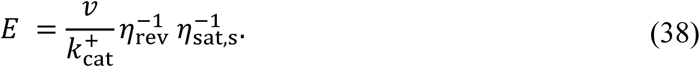

With this equation, we defined Enzyme Cost (EC) as the product of the enzyme demand and the enzyme molecular weight *w*:

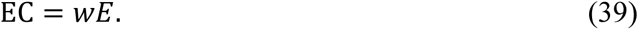

The enzyme molecular weights were retrieved from the UniProt database ^85^. We also defined pathway’s total EC as the sum of ECs in the pathway:

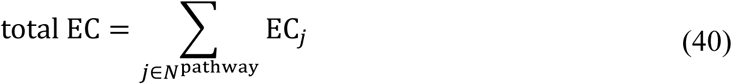

where 𝑁^pathway^ is reactions included in the pathway. We calculated ECs in glycolysis (Fig. 3A), gluconeogenesis (Fig. 3B), and the TCA cycle (Fig. 3C), which were the same as those used in FCC calculations, under the hepatic Δ_*r*_*G*′ landscape using metabolite concentrations and Δ_*f*_*G*′∘ values sampled for confidence interval calculation in GLEAM, and geometric mean of *K*_m_ values from the BRENDA database ^51^ obtained in FCC calculation (Table S5). For *K*_m_ with fewer than three data found in the BRENDA, we set their *c*/*K*_m_ values to the geometric mean of the pathway. *k*^+^ values were assumed to be the same within a pathway, and their effect can be ignored in this study because we only focused on the relative values of EC.

#### Minimization of total EC

Since EC is dependent on metabolite concentrations, we defined Enzyme Cost Minimum (ECM) as a theoretical condition where total EC is minimized with metabolite concentrations. We obtained ECM by solving the following non-linear optimization problem ^42^:

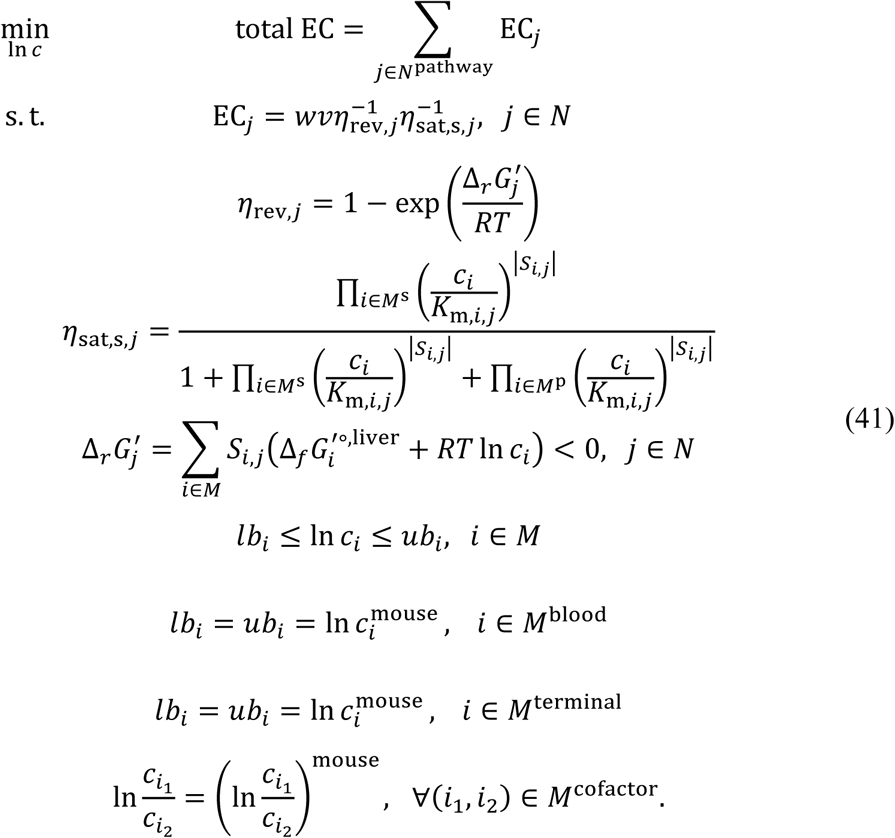

*w* and *v* are normalized by the minimum molecular weight and flux, respectively. The relative flux values for a pathway we analyzed were unique because the degree of freedom of these pathways’ steady flux is 1. ln *c*^mouse^ represents metabolite concentration in mouse *i.e.*, ln *c*^∗^ and ln *c*^𝑏∗^. *M*^blood^ represents blood metabolites, and *M*^terminal^ represents intracellular metabolites at the termini of each pathway: Glucose and Pyr for glycolysis and gluconeogenesis, and AcCoA, OAA, and Mal for the TCA cycle. By fixing these concentrations and cofactor ratios to those in mouse liver, the total Gibbs free energy change across the entire pathway in ECM is consistent with that in the mouse liver. ECM was calculated by MATLAB function fmincon with termination tolerance on the first-order optimality of OptimalityTolerance = 10^-7^, tolerance on the constraint violation of ConstraintTolerance = 10^-9^, maximum number of function evaluations of MaxFunctionEvaluations = 1000000, and maximum number of iterations of MaxIterations = 100000. The optimization was performed 100 times for a single problem, and the solution minimizing the objective function was selected as ECM.

#### Calculation of MEG

To quantify the efficiency of metabolic switching in terms of metabolite concentration changes, we defined the Metabolite Equilibrium Gap (MEG) as the minimum concentration changes required to reverse the direction of metabolism:

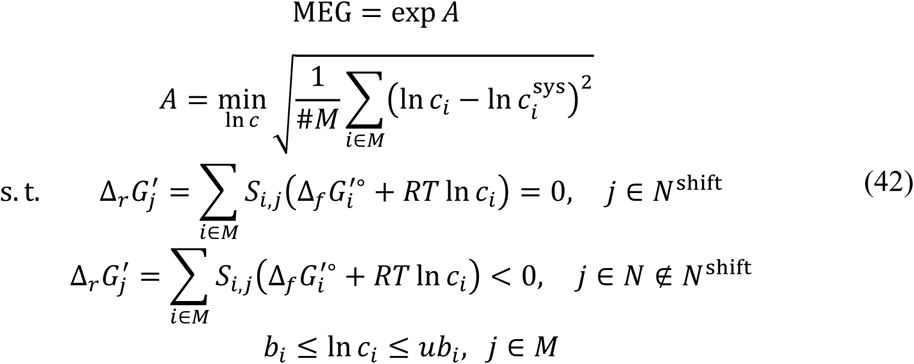

where sys represents a value in a target system corresponding to the liver or ECM in this study. 𝑁^shift^ is the reactions that must proceed in the reverse direction when switching metabolisms; it includes reversible reactions in glycolysis and gluconeogenesis, and all reactions in the TCA cycle. Δ_*f*_*G*′∘ values used in MEG calculation were the same as those in the liver (and also the same as those in ECM). MEG was calculated using Gurobi ^72^ MATLAB API with dual feasibility tolerance of OptimalityTol = 10^-9^ and primal feasibility tolerance of FeasibilityTol = 10^-9^.

### QUANTIFICATION AND STATISTICAL ANALYSIS

Statistical tests, optimizations, computational samplings, and calculations of Δ_*r*_*G*′, FCC, and EC were performed using MATLAB 2021b (The MathWorks Inc.) and Gurobi (version 10.0.3) ^72^. Database handlings were performed using Python (version 3.10. All details of statistical tests are described in the Results section. The MATLAB package Plot Groups of Stacked Bars ^86^ was used to create the figures.

## GLOSSARY

| Metabolite abbreviations | Metabolite names |
| --- | --- |
| G6P | Glucose 6-phosphate |
| F6P | Fructose 6-phosphate |
| F1,6BP | Fructose 1,6-bisphosphate |
| ADP | ADP |
| ATP | ATP |
| DHAP | Glycerone phosphate |
| GAP | Glyceraldehyde 3-phosphate |
| NAD | NAD<sup>+</sup> |
| NADH | NADH |
| Pi | Orthophosphate |
| 3PG | 3-Phospho-D-glycerate |
| 2PG | 2-Phospho-D-glycerate |
| PEP | Phosphoenolpyruvate |
| H<sub>2</sub>O | H<sub>2</sub>O |
| G1P | Glucose 1-phosphate |
| G3P | sn-Glycerol 3-phosphate |
| Pyr | Pyruvate |
| CoA | Coenzyme A |
| CO<sub>2</sub> | CO<sub>2</sub> |
| AcCoA | Acetyl-CoA |

| OAA | Oxaloacetate |
| --- | --- |
| Cit | Citrate |
| AKG | alpha-Ketoglutaric acid |
| SucCoA | Succinyl-CoA |
| Suc | Succinate |
| GDP | GDP |
| GTP | GTP |
| Fum | Fumarate |
| FAD | FAD |
| FADH2 | FADH2 |
| Mal | Malate |
| Glu | Glutamate |
| NH3 | Ammonia |
| Ala | L-Alanine |
| Lac | Lactate |
| HCO3 | HCO<sub>3</sub><sup>-</sup> |
| NADP | NADP<sup>+</sup> |
| NADPH | NADPH |
| Reaction abbreviations | Reaction names |
| GLUT | Glucose transporter |
| GCK | Glucokinase |
| HK | Hexokinase |
| GPI | Glucose-6-phosphate isomerase |
| PFK | Phosphofructokinase |
| ALDO | Aldolase |
| TPI | Triosephosphate isomerase |
| GAPDH:PGK | Glyceraldehyde-3-phosphate dehydrogenase and<br>Phosphoglycerate kinase |
| PGAM | Phosphoglycerate mutase |
| ENO | Enolase |
| PGM | Phosphoglucomutase |
| FBP | Fructose-1,6-bisphosphatase |
| G6PC | Glucose-6-phosphatase |
| GPD | Glycerol-3-phosphate dehydrogenase |
| PKLR | Pyruvate kinase |
| PDH | Pyruvate dehydrogenase |
| CS | Citrate synthase |
| ACO:IDH | Aconitase and Isocitrate dehydrogenase |
| OGDH | 2-Oxoglutarate dehydrogenase |
| LSC | Succinate-CoA ligase |
| SDH | Succinate dehydrogenase |
| FH | Fumarate hydratase |
| MDH | Malate dehydrogenase |
| GDH | Glutamate dehydrogenase |
| ALT | Alanine aminotransferase |
| LDH | Lactate dehydrogenase |
| MCT | Monocarboxylate transporter |
| PC | Pyruvate carboxylase |
| PCK | Phosphoenolpyruvate carboxykinase |
| ME | Malic enzyme |

## SUPPLEMENTAL INFORMATION

**Fig. S1:**
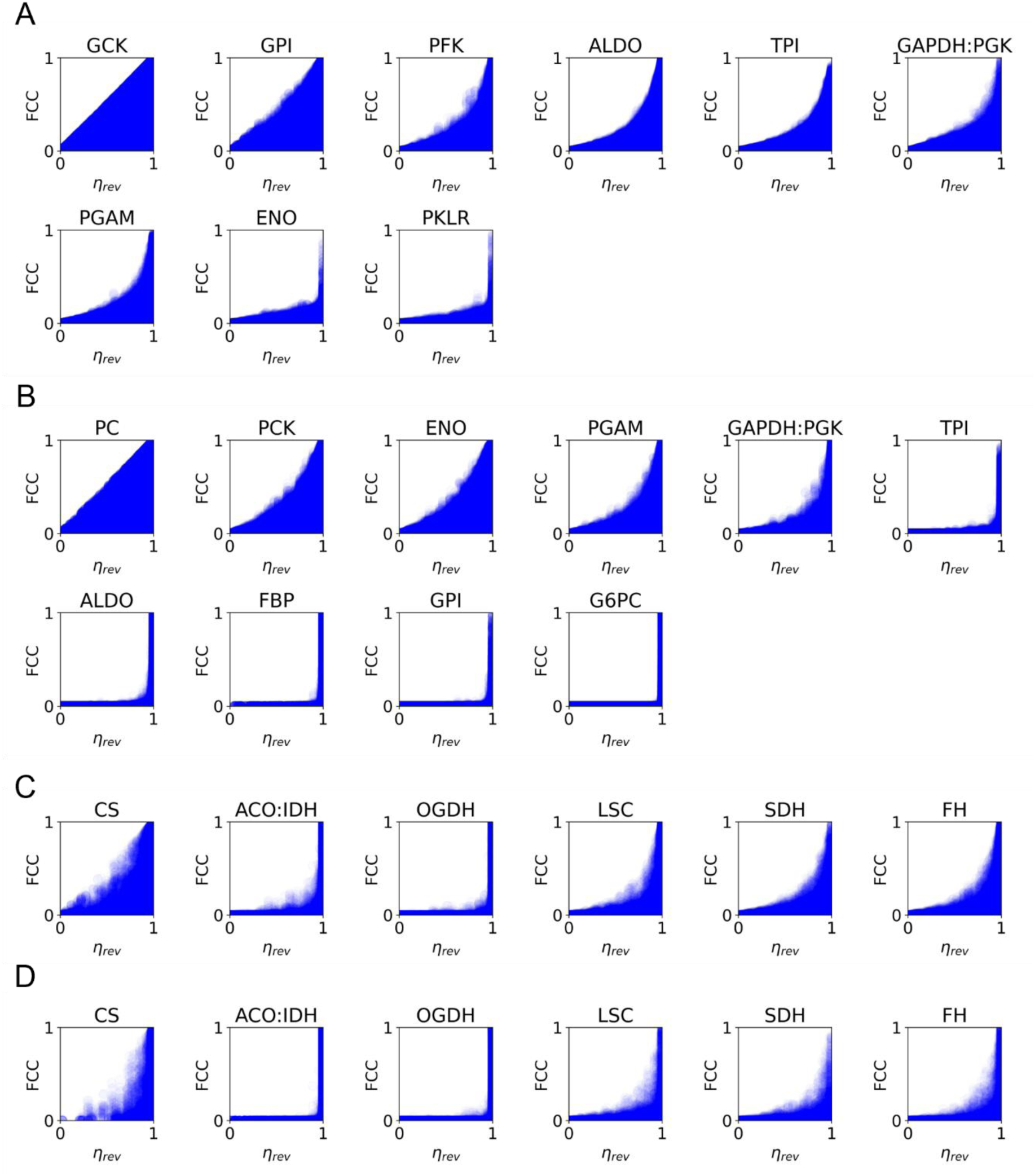
Relationship between 𝜼_𝐫𝐞𝐯_ and FCC for arbitrary Δ_*r*_*G*′ landscapes. (**A**-**D**) Arbitrary Δ_*r*_*G*′ landscapes were generated by the uniform sampling of intrasellar metabolite concentrations under the blood metabolite concentrations at *ad libitum* feeding and after 16-hour fasting (see Methods). The arbitrary Δ_*r*_*G*′ landscapes at *ad libitum* feeding were used for FCC calculation in glycolysis (**A**) and the TCA cycle (**C**), while those after 16-hour fasting were used for gluconeogenesis (**B**), and the TCA cycle (**D**). Plotted data represents each sampled set of metabolite concentrations and Δ_*f*_*G*′∘, and *K*_m_ values.

**Fig. S2:**
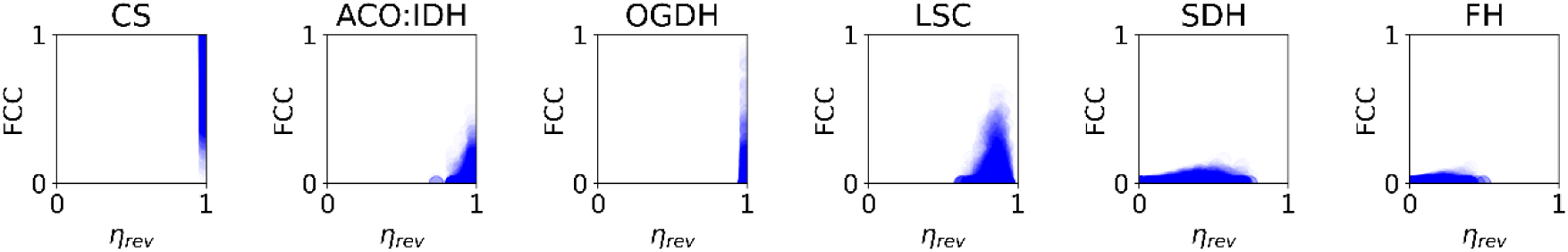
Relationship between 𝜼_𝐫𝐞𝐯_ and FCC in the TCA cycle in the mouse liver after 16-hour fasting. Plotted data represents each set of metabolite concentrations and Δ_*f*_*G*′∘ values sampled for the calculation of confidence intervals, and *K*_m_ values sampled based on BRENDA database (see Methods).

**Fig. S3:**
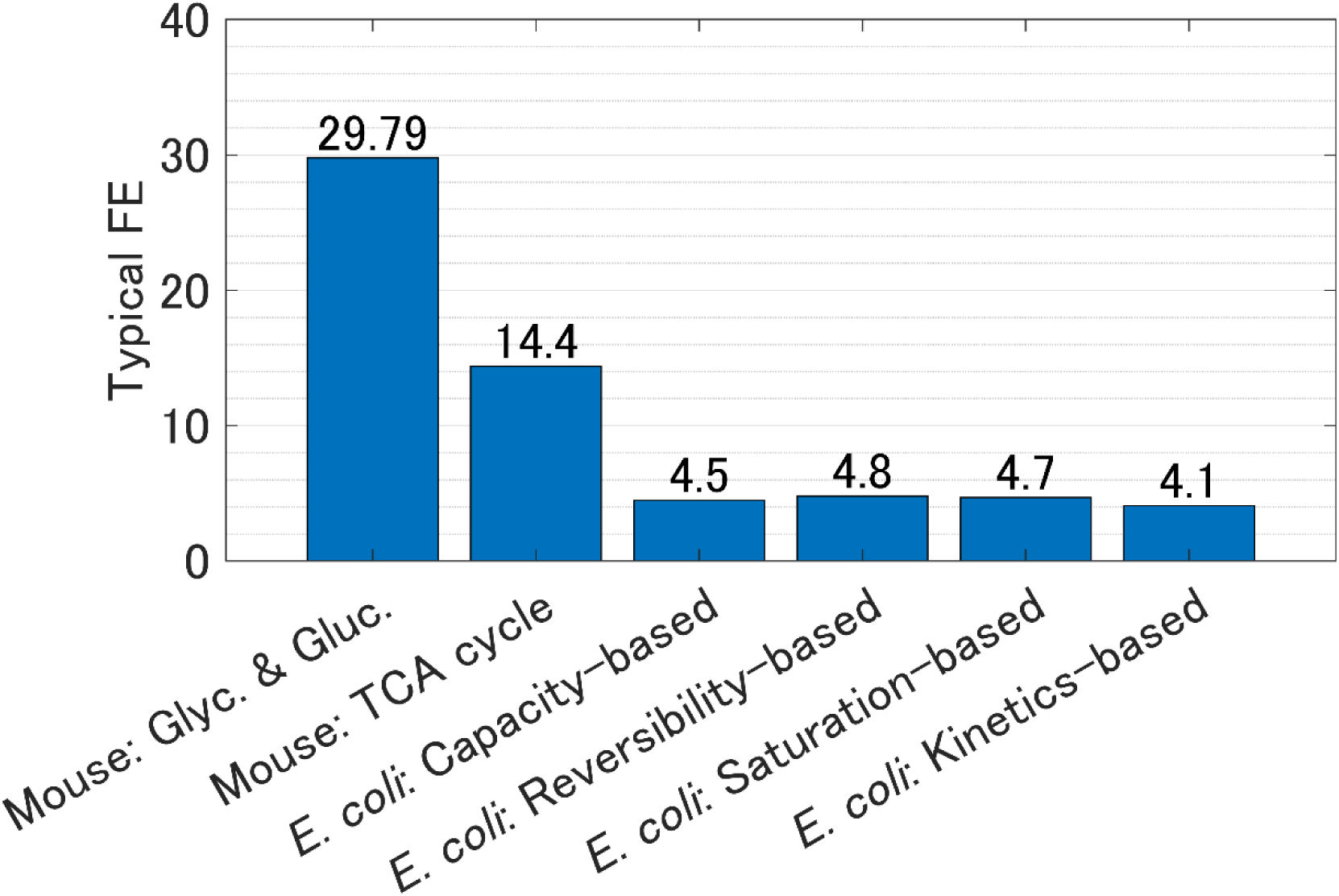
Typical fold error of ECM’s prediction compared to metabolite concentrations in the mouse liver and *E. coli*. Glyc. and Gluc. means glycolysis and gluconeogenesis, respectively. Minimizations of total EC in *E. coli* were conducted for the metabolic network, which consists of glycolysis, the pentose phosphate pathway, and the TCA cycle, assuming four different enzymatic rate laws (see Methods).

**Fig. S4:**
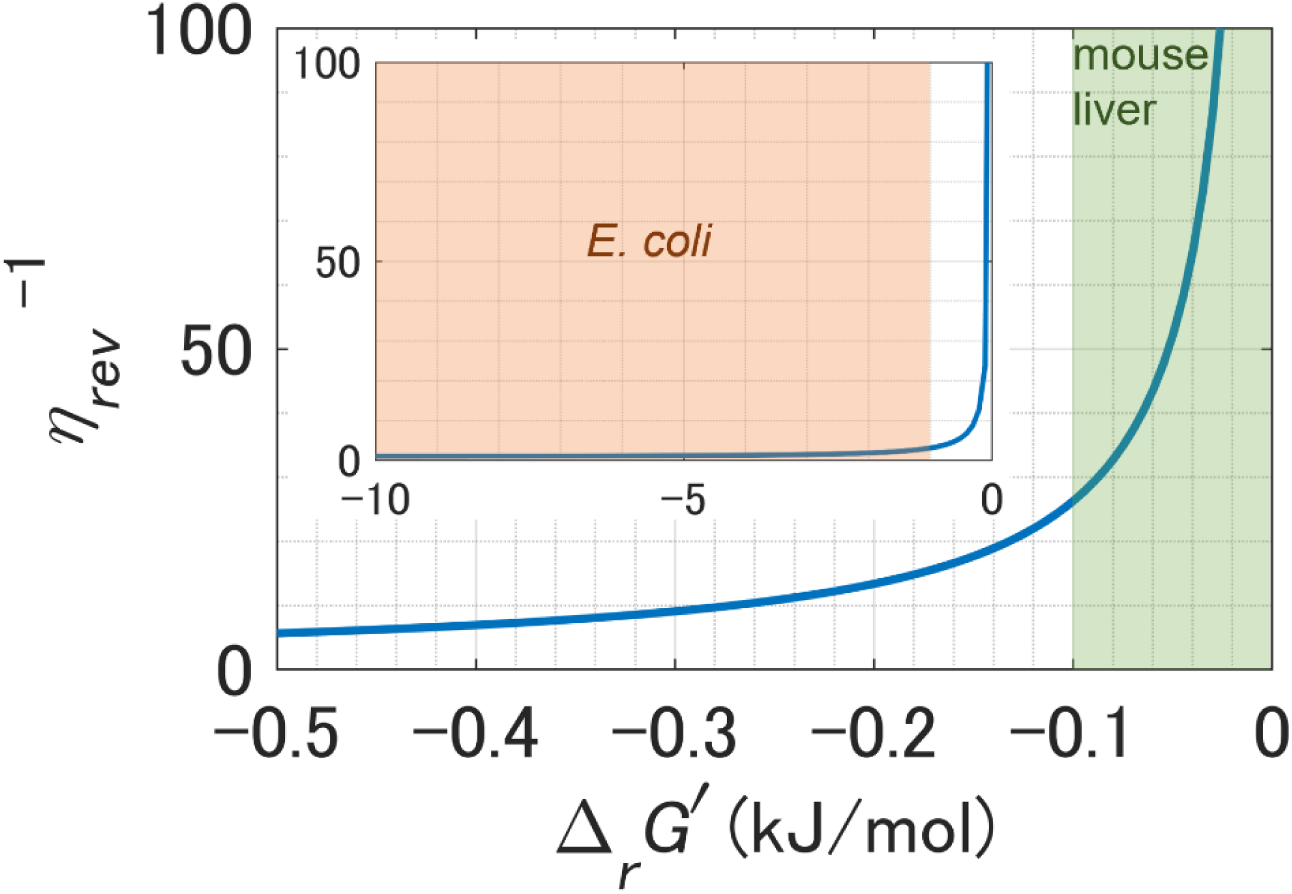
Correspondence between. Δ_*r*_*G*′ **and** 𝜼^−𝟏^ Orange and green area depict the range of Δ_*r*_*G*′ that most reversible reactions in glycolysis exhibited in *E. coli* ^26,60^ and mouse liver, respectively.

**Fig. S5:**
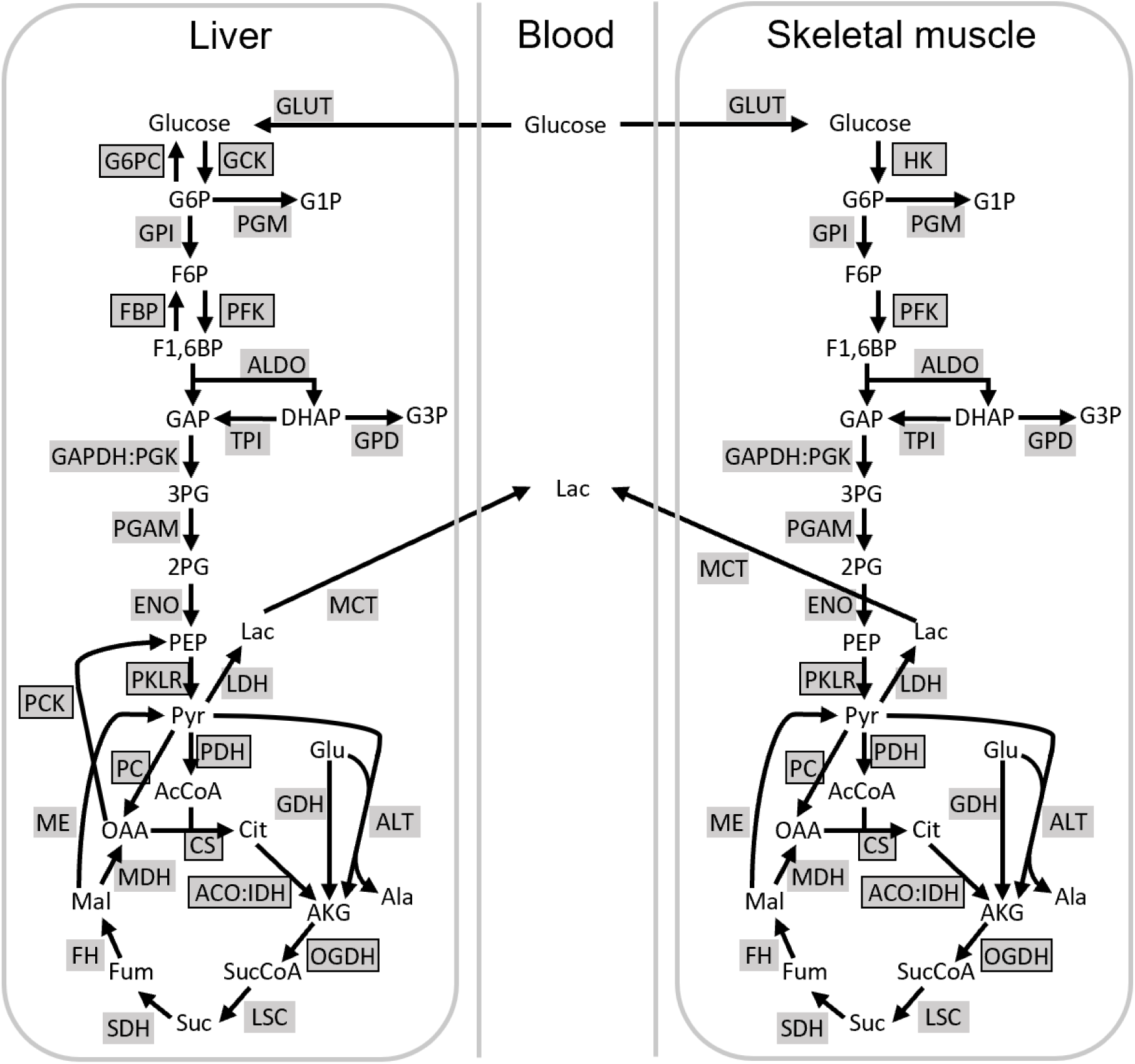
A whole metabolic network used to estimate Δ_*r*_*G*′ landscape. Shaded texts represent reaction names; boxed reactions are irreversible, while unboxed reactions are reversible. Unshaded texts represent metabolites. Although the network consists of liver, blood, and skeletal muscle to improve the accuracy of Δ_*f*_*G*′∘ estimation, only the Δ_*r*_*G*′ landscape of reactions and transporters in the liver were discussed in this study because the concentration of unmeasured major metabolites, such as glucose in skeletal muscle, cannot be estimated with sufficient accuracy due to a lack of reactions.

**Fig. S6:**
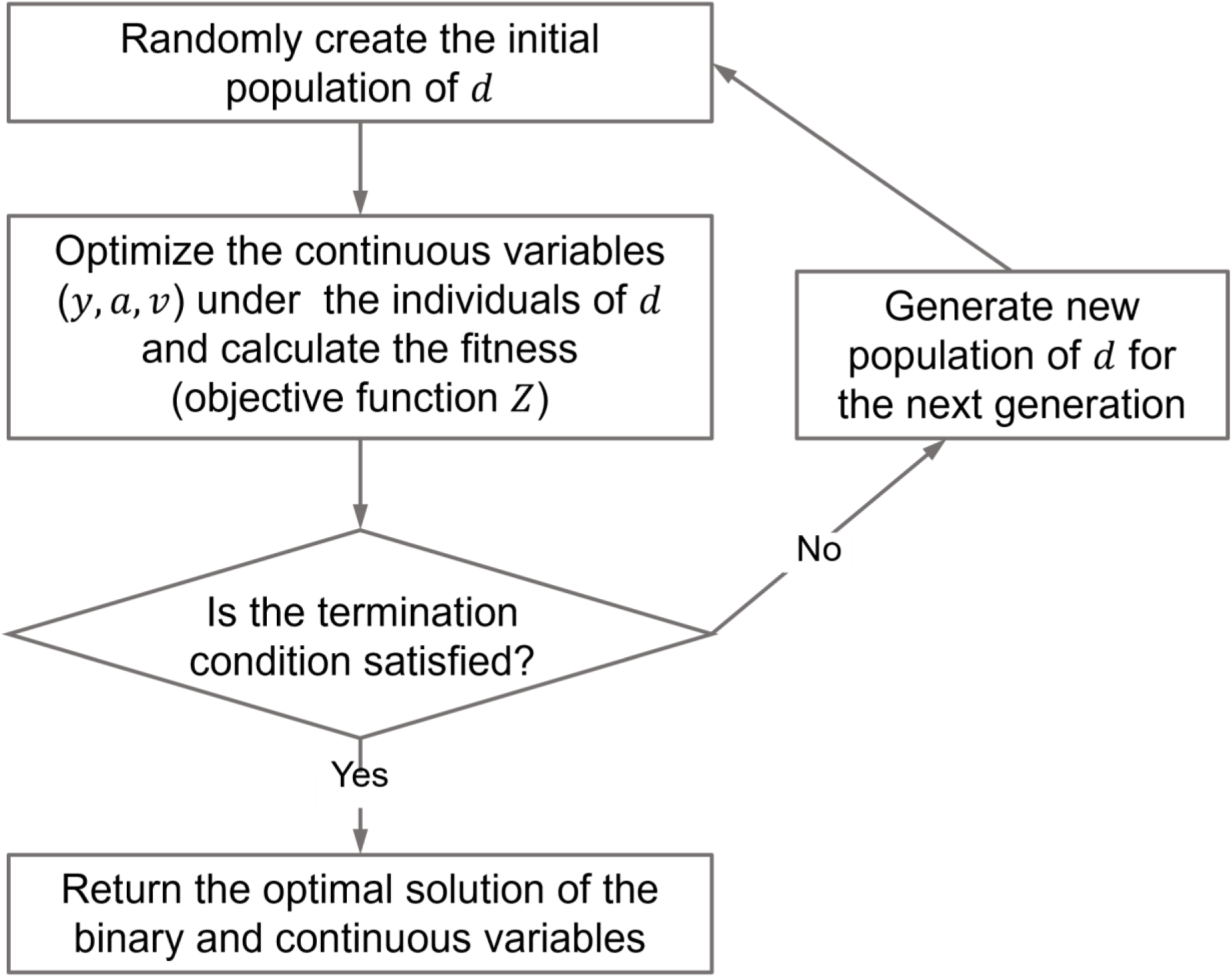
Flowchart of the bilevel optimization problem in GLEAM. The optimization problems for the continuous and binary variables are treated as the lower-level and upper-level problems, respectively. The optimal solution of GLEAM was selected after solving this bilevel optimization 10 times.

## SUPPLEMENTAL TABLES

**Table S1: Metabolomic and 𝚫_𝒇_𝑮′∘ data used as input for GLEAM (Excel file)**

This file includes the input metabolomic and Δ_*f*_*G*′∘ data for GLEAM (logarithmic mean of metabolome concentrations, Δ_*f*_*G*′∘ values, and variance-covariance matrix of metabolite concentrations and Δ_*f*_*G*′∘s). The metabolite concentrations were measured form from WT and *ob*/*ob* mice in previous studies ^8,66^, and Δ_*f*_*G*′∘ data were retrieved from eQuilibrator database ^55^. NaN represents unmeasured metabolite. [h], [m], and [b] indicate hepatic, myocytic, and blood compartment, respectively. The logarithmic mean of metabolite concentrations shown in this file is converted from a logarithmic scale to a linear scale.

**Table S2: Thermodynamically consistent metabolite concentrations, 𝚫_𝒇_𝑮′∘, and Δ_*r*_*G*′ estimated by GLEAM (Excel file)**

This file includes the thermodynamically consistent metabolite concentrations, Δ_*f*_*G*′∘, and Δ_*r*_*G*′ values in mouse liver and blood estimated by GLEAM. For glucose and lactate, hepatic and blood metabolites are distinguished by the [h] and [b], respectively.

**Table S3: Metabolites and reactions in the metabolic network for glucose metabolism in mice (Excel file)**

This file includes definitions of metabolite abbreviations, reaction abbreviations, and the metabolic network *i.e.*, the stoichiometric matrix for glucose metabolism used in this study. In the Reactions and Stoichiometric matrix sheet, [h], [m], and [b] indicate hepatic, myocytic, and blood compartments, respectively.

**Table S4:**
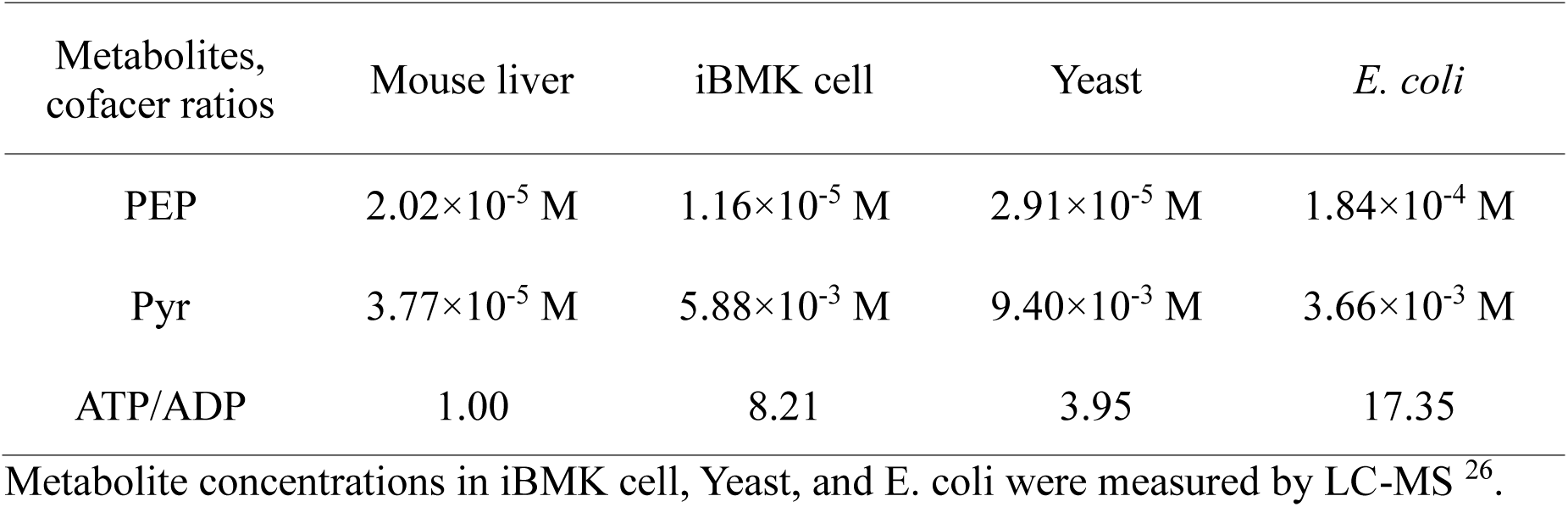
Metabolite concentrations and a cofactor ratio involved in PKLR between mouse liver and other cell types.

| Metabolites,<br>cofactor ratios | Mouse liver | iBMK cell | Yeast | *E. coli* |
| --- | --- | --- | --- | --- |
| PEP | $2.02 \times 10^{-5}$ M | $1.16 \times 10^{-5}$ M | $2.91 \times 10^{-5}$ M | $1.84 \times 10^{-4}$ M |
| Pyr | $3.77 \times 10^{-5}$ M | $5.88 \times 10^{-3}$ M | $9.40 \times 10^{-3}$ M | $3.66 \times 10^{-3}$ M |
| ATP/ADP | 1.00 | 8.21 | 3.95 | 17.35 |

Metabolite concentrations in iBMK cell, Yeast, and *E. coli* were measured by LC-MS<sup>26</sup>.

**Table S5: EC calculation in mouse liver and ECM (Excel file)**

This file includes EC values within glycolysis, gluconeogenesis, and the TCA cycle in mouse liver and EC, metabolite concentrations in ECM, and geometric mean of Michaelis-Menten constant retrieved from BRENDA database ^51^ used for EC calculation. In EC calculation, metabolite concentrations at *ad libitum* feeding were used for glycolysis and the TCA cycle, and those after 16-hour fasting were used for gluconeogenesis and the TCA cycle. EC is product of normalized molecular weight, 1/*η*_rev_, and 1/*η*_sat,s_.

**Table S6: Metabolite concentrations where the reactions required to reverse for metabolic switching are at equilibrium, as determined by MEG calculation (Excel file)** This file includes the metabolite concentrations that are the optimal solution to the optimization problem in MEG calculation for glycolysis, gluconeogenesis, and the TCA cycle. At these concentrations, the reactions required to reverse for directional switching of the pathways are at equilibrium with the minimum concentration changes from mouse liver or ECM.

**Table S7:** Upper and lower bounds of metabolite concentrations and cofactor ratios.

| Metabolites,<br>cofactor ratios | Lower bounds | Upper bounds | References |
| --- | --- | --- | --- |
| Intracellular glucose* | $1.0 \times 10^{-3}$ M | $1.5 \times 10^{-2}$ M | 87 |
| Hepatic Pi | $3.0 \times 10^{-3}$ M | $4.3 \times 10^{-3}$ M | 88 |
| Myocytic Pi | $3.0 \times 10^{-3}$ M | $9.0 \times 10^{-3}$ M | 88,89 |
| CO<sub>2</sub>, HCO<sub>3</sub><sup>-</sup> | $1.5 \times 10^{-3}$ M | $2.5 \times 10^{-3}$ M | 90 |
| NH<sub>3</sub> | $2.0 \times 10^{-4}$ M | $1.0 \times 10^{-3}$ M | 91 |
| Other intracellular<br>metabolites* | $1.0 \times 10^{-7}$ M | $1.0 \times 10^{-2}$ M | 26 |
| Blood metabolites | $1.0 \times 10^{-7}$ M | $1.0 \times 10^{-1}$ M | 92 |
| ATP/ADP | 1 | 10 | 89,93,94 |
| NAD<sup>+</sup>/NADH | 3 | 10 | 95 |
| GTP/GDP | 1 | 10 | 96 |
| NADP<sup>+</sup>/NADPH | 0.05 | 0.2 | 97 |

\*: The upper bounds of intracellular glucose, G6P, F6P, F1,6BP, DHAP, GAP, 3PG, 2PG,
PEP, G1P, G3P, and Lac in *ob/ob* mouse are $1.0 \times 10^{-1}$ M.

